# Epigenetic and transcriptional consequences of chemically induced transposon mobilization in the endosperm

**DOI:** 10.1101/2024.03.19.585643

**Authors:** Gerardo del Toro-de León, Joram van Boven, Juan Santos-González, Wen-Biao Jiao, Korbinian Schneeberger, Claudia Köhler

## Abstract

Genomic imprinting, an epigenetic phenomenon leading to parent-of-origin-specific gene expression, has independently evolved in the endosperm of flowering plants and the placenta of mammals—tissues crucial for nurturing embryos. While transposable elements (TEs) frequently colocalize with imprinted genes and are implicated in imprinting establishment, direct investigations of the impact of *de novo* TE transposition on genomic imprinting remain scarce. In this study, we explored the effects of chemically induced transposition of the Copia element *ONSEN* on genomic imprinting in *Arabidopsis thaliana*. Through the combination of chemical TE mobilization and doubled haploid induction, we generated a line with 40 new *ONSEN* copies. Our findings reveal a preferential targeting of maternally expressed genes (MEGs) for transposition, aligning with the colocalization of H2A.Z and H3K27me3 in MEGs— both previously identified as promoters of *ONSEN* insertions. Additionally, we demonstrate that chemically-induced DNA hypomethylation induces global transcriptional deregulation in the endosperm, leading to the breakdown of MEG imprinting. This study provides insights into the consequences of chemically induced TE remobilization in the endosperm, underscoring the need for cautious interpretation of the connection between TEs and genomic imprinting.

## INTRODUCTION

Transposable Elements (TEs) are genomic parasites that play essential and diverse roles in all living organisms. TEs can replicate and jump within their host genome, accounting substantially to the variation in genome size. Active transposition can result in gene disruption, chromosomal rearrangements, and heterochromatin expansion (1). Therefore, silencing mechanisms have evolved to prevent TEs from mobilizing, including the RNA-directed DNA methylation (RdDM) pathway in plants (2). Insertions of TEs can affect the expression of adjacent genes, either by silencing mechanisms spreading into gene flanking regions or by the TEs contributing regulatory elements impacting gene expression (3).

These mechanisms are particularly relevant in the endosperm, a seed tissue required to support embryo growth in flowering plants (4). The endosperm forms after double fertilization by the fusion of the central cell within the female gametophyte with one of the two sperm cells. The other sperm cell fertilizes the egg cell, giving rise to the embryo (5). The DNA glycosylase DEMETER (DME) is active in the central cell and removes methylated cytosines from small TEs, causing neighboring genes to be expressed (6). Since DME is not active in sperm, TEs remain methylated and flanking genes silenced. This mechanism establishes an epigenetic asymmetry of the parental genomes, resulting in parent-of-origin specific gene expression. This phenomenon is commonly referred to as genomic imprinting (4, 6, 7). Removal of DNA methylation in the central cell can also cause the recruitment of the Polycomb Repressive Complex2 (PRC2), leading to the establishment of the repressive trimethylation of histone H3 at lysine 27 (H3K27me3) (8–10) on maternal alleles. In consequence, the maternal alleles of the affected genes are silenced, while the paternal alleles are expressed (8, 10) .

Genomic imprinting has evolved independently in the embryo-nurturing tissues of flowering plants and mammals, evidencing a key role in reproduction (11). TEs often overlap with imprinted genes in both plants and mammals, leading to the hypothesis that TEs are responsible for establishing imprinting (12–15). Nevertheless, not all imprinted genes colocalize with TEs, and not all genes that overlap with TEs are imprinted (13). This disparity in observations highlights a gap in our understanding of the epigenetic regulation and evolutionary aspects of genomic imprinting. Therefore, studies addressing a direct and immediate effect on TE mobilization on genomic imprinting are relevant to close this knowledge gap.

Recent efforts to artificially mobilize TEs provide an opportunity to study the effects of transposition on imprinted gene expression on a laboratory time scale (16, 17). By combining the non-methylable cytosine analog zebularine with the Polymerase II inhibitor α-amanitin, transposition of the heat-responsive Copia element *ONSEN* could be successfully induced after heat treatment. Remobilized *ONSEN* elements have various effects on the transcriptional regulation of genes that are either directly targeted by *ONSEN* or flanking the insertion site, demonstrating the power of TEs to impact gene regulation (17).

Here, we *de novo* mobilized *ONSEN* and tested the effect on gene regulation in the endosperm of Arabidopsis. We found that imprinted maternally expressed genes (MEGs) were preferentially targeted by new *ONSEN* insertions, correlating with the enrichment of H3K27me3 and the histone variant H2A.Z that were previously found to promote *ONSEN* insertions (17, 18). These data suggest that the epigenetic landscape of MEGs are hotspots for new *ONSEN* insertions, implicating that imprinting can precede TE insertion. Furthermore, we found that new *ONSEN* insertions in the endosperm had low levels of CG and CHH methylation, but were marked by the heterochromatic H3K9me2 modification, suggesting that H3K9me2 is the primary silencing signal for new *ONSEN* elements in the endosperm. Finally, we observed a global breakdown of MEG imprinting, which we account to the combined effect of global and persistent DNA hypomethylation and the activation of transcription factors (TFs), inducing a global deregulation of gene expression in endosperm.

## MATERIAL AND METHODS

### Plant material and growth conditions

TEmob was generated in *Arabidopsis thaliana* using the previously reported endosperm specific INTACT line (isolation of nuclei tagged in specific cell types) (10) expressing *PHE1::NTF* and *PHE1::BirA* in the Col-0 accession and the *delayed dehiscence 2* (*dde2*) mutant background (19). The *dde2* mutation was segregated out during haploid induction. The INTACT line was used as wild-type (WT) reference for all comparisons. The *pistillata-1* (*pi-1*) mutant (20) was obtained from NASC. Seeds were surface sterilized with 70% ethanol and washed three times with sterile water. Sterilized seeds were sown on ½MS plates containing 0.5% sucrose and 0.8% agar, stratified for 2 days at 4°C and germinated under long-day conditions (16h light/8h darkness) at 21°C. Seedlings were transferred to soil after 10–12 days and grown in phytotrons under long day conditions (day 21°C, night 20°C 70% humidity, 150µE light intensity).

All endosperm datasets generated in this study (RNAseq, WGBS and CUT&Tag) were obtained from INTACT purified endosperm nuclei from Col × L*er* (L) reciprocal crosses at 4 days after pollination (DAP) using the INTACT line (referred as C) or TEmob (T3, referred as Cm) genetic background. To facilitate the crosses, we used the male sterile *pi-1* mutant in L*er* background as female parent pollinated with the INTACT lines. For the reciprocal cross TEmob and WT INTACT lines were emasculated and pollinated with L*er* wild-type pollen.

### Generation of the TEmob line

Chemical treatment for *ONSEN* mobilization was performed following the previously published protocol reported (16). In brief, seeds were sown on ½MS agar plates containing 5, 44µM α-amanitin (Sigma Cat. No. A2263) and 40µM zebularine (Sigma Cat. No. Z4775) stratified for 2 days at 4°C and germinated under long-day conditions (16h light/8h darkness) at 21°C. After 9 days, the plates were incubated at 4°C degrees for 24h and subsequently at 37°C in the dark for 24h for heat shock induction (HS). *ONSEN* copy number was estimated by qPCR in treated plants (T0) and their progeny (T1) (**Supplementary Table S1**). Plants with the highest *ONSEN* copy numbers were selected and crossed with homozygous *cenh3 CENH3- TAILSWAP* plants (21). Offspring T2 haploids were grown and treated with colchicine. Ploidy was measured on a CyFlow Ploidy Analyser (Sysmex). A single diploid line was identified (TEmob) in the following generation (T3).

### Whole genome Illumina sequencing and analysis

Genomic DNA for short-read sequencing was extracted from 5-week-old leaves using the MagJET™ Plant Genomic DNA Kit (Thermo Scientific, Cat. No. K2761). DNA libraries were prepared using the NEBNext® Ultra™ II DNA Library Prep Kit for Illumina® (New England Biolabs Cat. No. E7645) and the NEBNext® Multiplex Oligos for Illumina® (New England Biolabs Cat. No. E7335S). Size selection was aimed at 300-400 bp fragments. Sequencing was performed at Novogene on a HiSeq X platform in 150-bp paired-end (PE) mode.

### Whole genome PacBio sequencing

For high quality DNA isolation, 2g of flash-frozen 12 day-old seedlings were ground in liquid Nitrogen and DNA was extracted using 8mL of CTAB buffer (22). DNA purification was performed with 1x volume of Phenol:Chloroform:Isoamyl Alcohol (25:24:1, pH 8.0) and precipitated with 0.7x volume of isopropanol. The pellet was washed with 75% ethanol and gently resuspended in sterile water and treated with RNAse A (Thermo Scientific™ EN0531) and Ribonuclease T1 (Thermo Scientific™ EN0541) for 30 minutes at 37°C. The enzymes were extracted by adding 1x volume of Phenol:Chloroform:Isoamyl Alcohol (25:24:1, pH 8.0) and remaining phenol was removed with 1.5x volume of Chloroform. The DNA was precipitated for 30 min at -20°C with 1/10 volume of 3M Sodium acetate, pH 5.3 and 2.5 volume of 100% ethanol and resuspended in sterile water. Final DNA purification was performed with the Zymogen Genomic DNA Clean and Concentrator-10 column (Zymo Research, Irvine). Libraries were prepared using SMRTbell Template Prep Kit 1.0 and sequenced on a PacBio Sequel platform at Novogene.

### TEmob chromosome-level assembly and annotation of TEmob

PacBio short (<50 bp) and low quality (QV < 80) reads were filtered out using the SMRTLink5 package. PacBio long reads were *de novo* assembled with Flye tool (23) and polished with the Arrow tool from the SMRTLink5 package after remapping with the pbmm2 (minimap2) tool. The resulting assembly was further corrected to remove small-scale assembly errors with bcftools based on the alignments generated with the Illumina short reads using BWA (24). The final chromosome-level assembly was generated based on the synteny alignment with the Col-0 TAIR10 reference genome using MUMmer4 and the Perl scripts described by (25). Gene annotation was performed using the tool liftoff with the TAIR10 reference annotation. TEs were annotated with RepeatMasker (http://www.repeatmasker.org).

### Analysis of new *ONSEN* insertions

TE annotations from the reference Col-0 wild type and TEmob were compared and significant differences in the number of members per family were only observed for *ATCOPIA78* (*ONSEN*). Manual curation was done by comparing each *ONSEN* annotation between the wild-type reference and TEmob. Analysis of epigenetic modifications was performed by analyzing 100bp upstream and downstream of the new insertions and targeted genes with the map feature of bedtools v2.30.0 (26). Plots and statistical analysis were done in R.

### Analysis of H3K27me3 and H2A.Z leaf datasets

We made use of publicly available datasets for H3K27me3 (GSE66585) and H2A.Z (GSE123263) from leaf tissues (10, 27). Data processing was performed as previously described by (10). Analysis, plots and statistical analysis were done with bedtools v2.30.0 and R (26).

### Aerial tissue whole-genome bisulfite sequencing and analysis

DNA was extracted from 100 mg of ground flash-frozen aerial tissue pooled from three 4- week-old TEmob (T4) plants using the MagMAX™ Plant DNA Isolation Kit in biological duplicates. Libraries were prepared with the Accel-NGS Methyl-Seq DNA Library Kit (Swift Cat No. 30096), and the sequencing was performed at Novogene on a NovaSeq 6000 platform in 150-bp PE mode. For the wild-type reference, we used the WGBS dataset from GSE156597 previously generated in our group using comparable specifications (3 week-old aerial tissues) (28). 150-bp PE reads were trimmed by removing the first 5 bases from the 5’ end and the last 20 bases from the 3’ end. Reads were mapped to the Arabidopsis TAIR10 in PE mode (-- score_min L, 0, -0.6) genome using Bismark (29). Duplicated reads were eliminated and methylation levels for each condition were calculated by averaging the two biological replicates. Differentially methylated regions (DMRs) between TEmob and WT were defined using 50-bp windows across the genome as units. Cytosine positions with at least 6 informative mapped reads and 50bp windows with at least 4 cytosines were considered. Windows with differences in fractional methylation below the 1st decile (Fisher’s exact test, *P*-value < 0.01) were selected. DMRs with absolute methylation differences of 0.35 for CG, 0.20 for CHG and 0.15 for CHH (Fisher’s exact test, *P*-value < 0.01) were analyzed and merged if they occurred within 300 bp. Genomic features indicated as gene (gene-body), TEs, promoter (1Kb 5’ upstream) and intergenic regions were defined based on the current TAIR10 genome release and overlapping with DMRs were obtained using intersect feature of bedtools v2.30.0 (26).

### Endosperm RNA sequencing and analysis

A total of 250 mg of siliques at 4DAP were collected from TEmob Col (Cm) × L*er* (L) reciprocal crosses in three biological replicates. Tissue homogenization, nuclei purification, RNA extraction, and library preparation were performed as previously described (30, 31). Libraries were sequenced at Novogene on a HiSeq platform in 150-bp PE mode. For each replicate, the 150 bp reads were trimmed and mapped in single-end mode to the Arabidopsis (TAIR10) genome masked for rRNA genes using TopHat v2.1.0 (32) (parameters adjusted as -g 1 -a 10 -i 40 -I 5000 F 0 r 130). Mapped reads were counted in the genes using GFOLD (33). Differentially regulated genes were detected using DESeq2 (v. 1.42.0) (34) and R (v. 4.3.0). Comparisons were made against our previously published wild-type samples generated and processed as explained above (GSE119915). Only changes in expression with FDR of <0.01 and an absolute Log2 Fold Change (LFC) >= 1.5 were considered statistically significant. Parent-of-origin allele specific expression analysis and criteria to define imprinted genes was done as previously described (31, 35) following a stricter minimum threshold of 50 informative reads. To increase the statistical power to detect parentally biased genes, replicates were merged.

### Endosperm whole-genome bisulfite sequencing and analysis

A total of 500 mg of siliques at 4DAP were collected from wild-type Col (C) x L*er* (L) reciprocal crosses and from TEmob Col (Cm) × L*er* (L) reciprocal crosses in biological duplicates. Tissue homogenization and nuclei purification were performed as previously described (30, 31). DNA was extracted with the DNeasy® Plant Mini Kit (Qiagen Cat. No. 69104). Libraries were prepared with the Accel-NGS Methyl-Seq DNA Library Kit (Swift Cat No. 30096), and the sequencing was performed at Novogene on a NovaSeq 6000 platform in 150-bp PE mode. Bisulfite sequencing data processing and parent-of-origin methylation was performed as previously described (10). DMRs identification and analysis were performed as described for the aerial tissue samples above.

### Endosperm INTACT-CUT&Tag sequencing and analysis

A total of 250 mg of siliques at 4DAP were collected from wild-type Col (C) x L*er* (L) reciprocal crosses and from TEmob Col (Cm) × L*er* (L) reciprocal crosses in biological duplicates. We coupled our INTACT protocol for endosperm-nuclei purification with Cleavage Under Targets & Tagmentation (CUT&Tag) (36) for epigenomic profiling of H3K9me2 and H3K27me3. Tissue homogenization and nuclei purification were performed as previously described (30, 31) except for the replacement of MgCl2 by spermidine 0.5mM in the Honda Buffer to prevent pAG-Tn5 activation by residual MgCl2. We performed all steps with streptavidin-bound nuclei. After purification, nuclei were resuspended in Antibody150 buffer and from antibody incubation to tagmentation with pAG-Tn5 (EpiCypher Cat No. 15-1017) we followed the CUT&Tag Protocol v1.7 from CUTANA™. Primary antibody incubation was performed overnight at 4°C with gentle shaking using anti-H3K9me2 (Diagenode Cat. No. C15410060), Anti-Histone H3 (Sigma-Aldrich Cat. No. H9289) or anti-H3K27me3 (Cell Signaling Technology Cat. No. 9733T) antibodies. Secondary antibody incubation was performed with Guinea Pig anti-Rabbit IgG, (Antibodies-Online Cat. No. ABIN101961). Tagmentation was performed at 37°C for 1hr. PCR library amplification and post-PCR cleanup was performed as recommended by (37). We prepared unique index Nextera-compatible libraries with custom-designed index primers (38) amplified with Non-hot Start Phusion™ High-Fidelity DNA Polymerase (Thermo Scientific™ Cat No. F530S) for 14 cycles. Sequencing was performed on a NextSeq 1000 platform in 150- bp PE mode. PE reads were trimmed using the trim_galore tool and mapped to the Arabidopsis (TAIR10) genome using Bowtie2 (39) (parameters --no-unal --no-mixed --no-discordant --phred33 --local --very-sensitive-local). Mapped reads were sorted and indexed using SAMtools (40). Read coverage was estimated and normalized to 1x sequencing depth (reads per genome coverage, RPGC) using the bamCoverage from deepTools (41) (parameters -bs50 --effectiveGenomeSize 119481543 --normalizeUsing RPGC --skipNAs). Signal-to-noise ratio was normalized with H3 data by calculating the log2 ratio in 50 bins across the genome using bigwigCompare from deepTools. Parent-of-origin methylation was performed as previously described (10). For comparative purposes across samples data were standardized and normalized with a z-score transformation (42).

### Motif analysis

Annotation of Transcription Factors and predicted motifs were obtained from the Arabidopsis transcription factor database (AtTFDB) and from the Plant Transcription Factor Database (PlantTFDB 5.0) (43, 44) (http://planttfdb.gao-lab.org/). 1kb and 3kb upstream sequences to the ATG start codon were obtained from TAIR (https://www.arabidopsis.org). Motif analysis was performed using the fimo tool (45). Plots and statistical analysis were performed in R.

## RESULTS

### TEmob- A double haploid line with 40 new *ONSEN* insertions

To test how *de novo* transposition influences imprinting in Arabidopsis, we followed a published protocol for chemically inducing *ONSEN* TE mobilization (16). We applied the treatment to a previously established endosperm-specific INTACT line that allows purification of endosperm nuclei for downstream experiments (10). To ensure homozygosity of remobilized TEs, we used lines with higher *ONSEN* copy number (T1) as pollen donor and crossed them to a *cenh3 CENH3-tailswap* haploid inducer line (21). Haploid progenies were identified (T2) and treated with colchicine to duplicate the genome. A single double haploid line was successfully recovered (T3) which we refer as TEmob hereafter (**Supplementary Figure S1**).

To accurately determine the location of newly inserted TEs in the TEmob genome, we performed whole-genome sequencing with long read technology from PacBio. TEmob genome assembly and annotation confirmed an increased number of *ONSEN* elements. In addition to the 8 full-length native *ONSEN* elements in the Col-0 genome, 42 new full-length *ATCOPIA78* (*AtONSENmob1* to *42*) elements were detected by RepeatMasker in TEmob, distributed in all five chromosomes (**Figure 1A, B, Supplementary Figure S2**). Two of the new *ONSEN* TEs (*AtONSENmob9* and *AtONSENmob28*) are likely native to Col-0 and were potentially misannotated in the TAIR10 assembly, in agreement with a recent report (46) (**Supplementary Table S2**). Thus, a total number of 40 new full-length *ONSEN* elements were detected in the TEmob genome (**Figure 1A, B, Supplementary Table S2**). Analyses of sequence polymorphisms between full-length native and new *ONSEN* elements in TEmob revealed that most *AtONSENmob* were derived from *AT1TE12295* (*AT1G11265*) and few from *AT1TE59755* (*AT1G48710*) (**Figure 1A, Supplementary Figure S3**). These copies contain heat responsive elements (HREs), making them prone to be activated by heat treatment (47). No other TE family was found to increase in copy number in TEmob, consistent with previous reports showing that the applied protocol specifically mobilizes the heat-responsive Copia-like *ONSEN* retrotransposon (16, 17).

**Figure 1.**
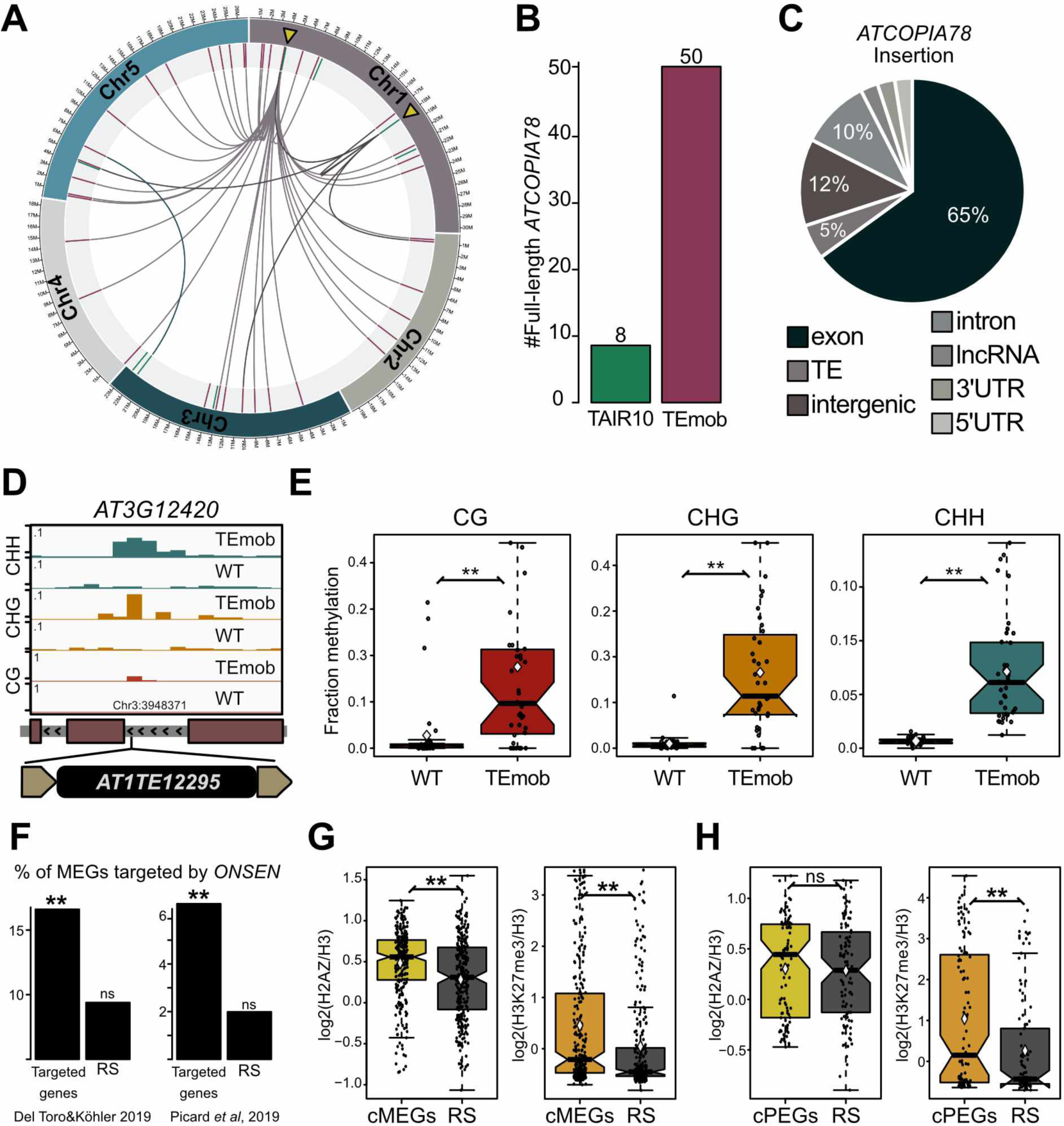
Characteristics of new full-length *ONSEN* insertions in TEmob. (**A**) New *ONSEN* insertions are distributed in the 5 chromosomes of *A. thaliana*. *AT1TE12295* and *ATITE59755* (yellow triangles) account for most of the insertions. (**B**) Total number of full-length *ONSEN* in TAIR10 and TEmob genomes. (**C**) Genomic context of new *ONSEN* insertions. (**D**) Genome browser example of a new insertion located in an intron of *ATG12420* showing enrichment of CG, CHG and CHH methylation in flanking insertion regions of TEmob in comparison to WT. (**E**) Leaf methylation levels in WT and TEmob in 100bp flanking insertion regions (upstream and downstream) of new *ONSEN* elements. (**F**) Percentage of MEGs targeted by *ONSEN* reported by Roquis *et al* 2021 in comparison to the percentage expected from a random sample (RS) of the same size. Two independent imprinting datasets depicted side by side. Significance was tested using a Hypergeometric test by comparing to the observed MEGs in each dataset. (**G**) H2A.Z (left) (Potok *et al* 2019) and H3K27me3 (right) (Moreno-Romero *et al* 2017) occupancy in core MEGs (cMEGs) and (**H**) core PEGs (cPEGs) in aerial tissues (cMEGs and cPEGs are genes reported as imprinted in at least two independent datasets, Supplementary Table S3). Wilcoxon test (E, G and H), \*\**P*-value <0.01; \**P*-value <0.05; ns, not significant.

To understand the potential effect of newly inserted *ONSEN* elements on neighboring gene expression, we performed whole-genome bisulfite sequencing (WGBS) on vegetative aerial tissues and analyzed changes in DNA methylation (DNAme) at the insertion regions. Of the new *ONSEN* elements, 65% were inserted into exons, consistent with previous observations of insertion preferences into exons for this family of retrotransposons, followed by intergenic regions and introns (**Figure 1C, Supplementary Table S2**). New *ONSEN* elements recruited high levels of DNAme in all sequence contexts to flanking regions of insertion sites (100bp upstream and downstream of the new insertions) (**Figure 1D, E**), revealing a strong transcriptional gene silencing (TGS) of newly transposed elements and spreading of DNAme into flanking regions. There was a minor increase of DNAme in the native full-length *ONSEN* copies, demonstrating reinforcement of silencing after TE mobilization (**Supplementary Figure S4**).

In summary, we established an *ONSEN*-mobilized double haploid line by combining chemical TE mobilization followed by maternal genome elimination and subsequent colchicine-induced diploidization (**Supplementary Figure S1**). The resulting TEmob line contains 40 new full-length *ONSEN* insertions that exhibited high CG and non-CG DNAme in aerial tissues at the flanking insertion sites, potentially influencing gene regulation in neighboring regions. The TEmob line was phenotypically indistinguishable from WT, as we did not observe any obvious differences in vegetative growth under normal growth conditions.

### New *ONSEN* insertions preferentially target MEGs

TEs frequently colocalize with imprinted genes (13, 48, 49), leading to the hypothesis that epigenetic regulation of TEs is connected to genomic imprinting. To study potential effects on genomic imprinting derived from new *ONSEN* copies in TEmob, we first analyzed the imprinting status of genes overlapping or being in the vicinity of new *ONSEN* elements prior *de novo* insertion. Of the 40 new insertions, only 11 were located within or in the vicinity (1kb up or downstream) of genes expressed in the endosperm and had sufficient read depth over single nucleotide polymorphisms (SNPs) in available imprinting datasets to distinguish parental alleles (13, 31, 35, 48, 50–52) (**Supplementary Table S3**). Interestingly, out of the 11 genes colocalizing with new *AtONSENmob* elements, 10 genes are imprinted or have parentally biased expression (**Supplementary Table S4**). In datasets of Col (C) and L*er* (L) accessions, *HDG3* (*AT2G32370*) is a paternally expressed gene (PEG), *UGT71C4* (*AT1G07250*), *AT1G46552*, *MCTP16* (*AT5G17980*) and *AT3G20975* are MEGs in CxL and *UGT71C4* and *AT3G20975* show significantly maternal biased expression in the reciprocal cross LxC. Two genes, *UGT71C4* and *MCTP16* also show significantly maternal biased expression in an independent study with Col-Cvi (CxV) accessions (52) (**Supplementary Table S4**). Further analysis of published imprinted expression datasets (13, 31, 35, 48, 50–52) revealed additional imprinting for *VTE* (*AT5G39220*) and *AT4G29580*, which are reported as MEGs and for *AT2G21930* as a PEG (**Supplementary Table S4**). Finally, *GSTF3* (*AT2G02930*) is not imprinted but shows significantly maternally biased expression in the CxV dataset (**Supplementary Table S4**) (52). Since imprinted genes are only a small subset of genes present in the endosperm ranging from 3 to 7% depending on the study (31, 52), we conclude that new *ONSEN* insertions in TEmob are overrepresented in this group of genes.

To test whether preferential insertion of *ONSEN* into imprinted genes is a general trend or specific to TEmob, we analyzed parent-of-origin expression of genes overlapping with previously reported *ONSEN* insertions (17) (**Supplementary Table S5**). From the 237 genes associated with new *ONSEN* insertions (17), 78 had sufficient read depth over SNPs in CxL (31) and 124 in CxV reciprocal crosses (52). Of those genes overlapping with new *ONSEN* insertions, 16.7% were identified as MEGs in CxL parent-of-origin expression data, which is significantly more than expected by chance (9%, *P*-value<0.004, Hypergeometric Test) (**Figure 1F and Supplementary Table S6**). Similarly, when comparing to the CxV dataset (52), 6.5% of *ONSEN* overlapping genes were MEGs, which is significantly more than expected (2.4%, *P*-value<0.003, Hypergeometric Test) (**Figure 1F and Supplementary Table S6)**. These observations support that imprinted genes and specifically MEGs are preferentially targeted by new *ONSEN* insertions.

It was reported that *ONSEN* preferentially targets silenced genes enriched for H3K27me3 and the histone variant H2A.Z (17, 18). We analyzed previously published epigenome profiles from aerial tissues (10, 27) to determine whether H2A.Z and H3K27me3 occupancy explains *ONSEN* preferential insertion into MEGs. First, we analyzed H2A.Z and H3K27me3 occupancy in genes associated with new *ONSEN* insertions in TEmob and found that indeed these regions were enriched for both H2A.Z and H3K27me3 (**Supplementary Figure S5**). We further tested whether imprinted genes were occupied by H2A.Z and H3K27me3 in aerial tissues and also found a substantial enrichment of H2A.Z and H3K27me3 over MEGs compared to randomly selected genes (**Figure 1G**). In contrast, PEGs were significantly enriched for H3K27me3 but not for H2A.Z (**Figure 1H**). Together, our data show that *de novo ONSEN* insertions preferentially target MEGs, possibly a consequence of MEGs being enriched for H2A.Z and H3K27me3 in aerial tissues.

### Recruitment of repressive epigenetic modifications in the endosperm by new *ONSEN* insertions

Since new *ONSEN* elements inserted into imprinted genes and recruited DNA methylation to flanking regions in vegetative aerial tissues (**Figure 1D, E**), we tested whether this would be reflected by DNA and histone methylation changes in TEmob endosperm. We generated endosperm-specific DNA methylation profiles at 4DAP in reciprocal crosses between L*er* (L) with either TEmob (Cm) or WT (C): CxL; LxC and CmxL; LxCm (also referred as C/CmxL and LxC/Cm). In contrast to vegetative aerial tissues, new *ONSEN* flanking sites did not contain significantly increased CGme and CHHme on maternal and paternal alleles in the endosperm (**Figure 2A, B**). Interestingly however, when TEmob was the maternal parent (CmxL), flanking regions of new *ONSEN* insertions showed a strong enrichment of CHGme, which was not observed when TEmob was the paternal parent (LxCm) (**Figure 2A, B**). CHG methylation generally co-occurs with H3K9me2 and in PEGs both marks colocalize with H3K27me3 to jointly repress maternal alleles ((10), **Supplementary Figure S6 and S7**). To test if new *ONSEN* insertions recruit heterochromatic marks in the endosperm, we generated H3K9me2 and H3K27me3 CUT&Tag epigenome profiles from 4DAP INTACT purified endosperm of reciprocal crosses between L*er* (L) with either TEmob (Cm) or WT (C): CxL; LxC and CmxL; LxCm. In agreement with the maternal inheritance of CHGme at flanking regions of new *ONSEN* insertions, we observed a significant enrichment of H3K9me2 when TEmob was the maternal parent in CmxL (**Figure 2C, Supplementary Figure S8A**). Though we did not detect CHGme at new *ONSEN* insertions when TEmob was paternally inherited, these regions were significantly enriched for H3K9me2, which was also reflected by increased H3K9me2 deposition on the entire targeted gene (**Figure 2B and C, Supplementary Figure S9A**). These observations suggest that H3K9me2 in the flanking regions of newly inserted *ONSEN* elements can be established independently of CHGme in the early endosperm. Maintenance of H3K9me2 depends on SUVH4/5/6, which are expressed in the endosperm (53). H3K9me2 acts in a reinforcing loop with non-CGme (54). However, H3K9me2 deposition was shown to occur independently of DNAme by sRNA-dependent recruitment of SUVH9 in the Arabidopsis embryo (55). Furthermore, SUVH9 (AT4G13460) is associated with H3K9me2 deposition independently of non-CGme enrichment in the endosperm of 3x seeds (53). We thus conclude that new *ONSEN* elements recruit H3K9me2 to their flanking regions that can be inherited and maintained in the endosperm. These observations also suggest that H3K9me2 at new *ONSEN* elements in the paternal genome can be deposited in a DNAme independent way.

**Figure 2.**
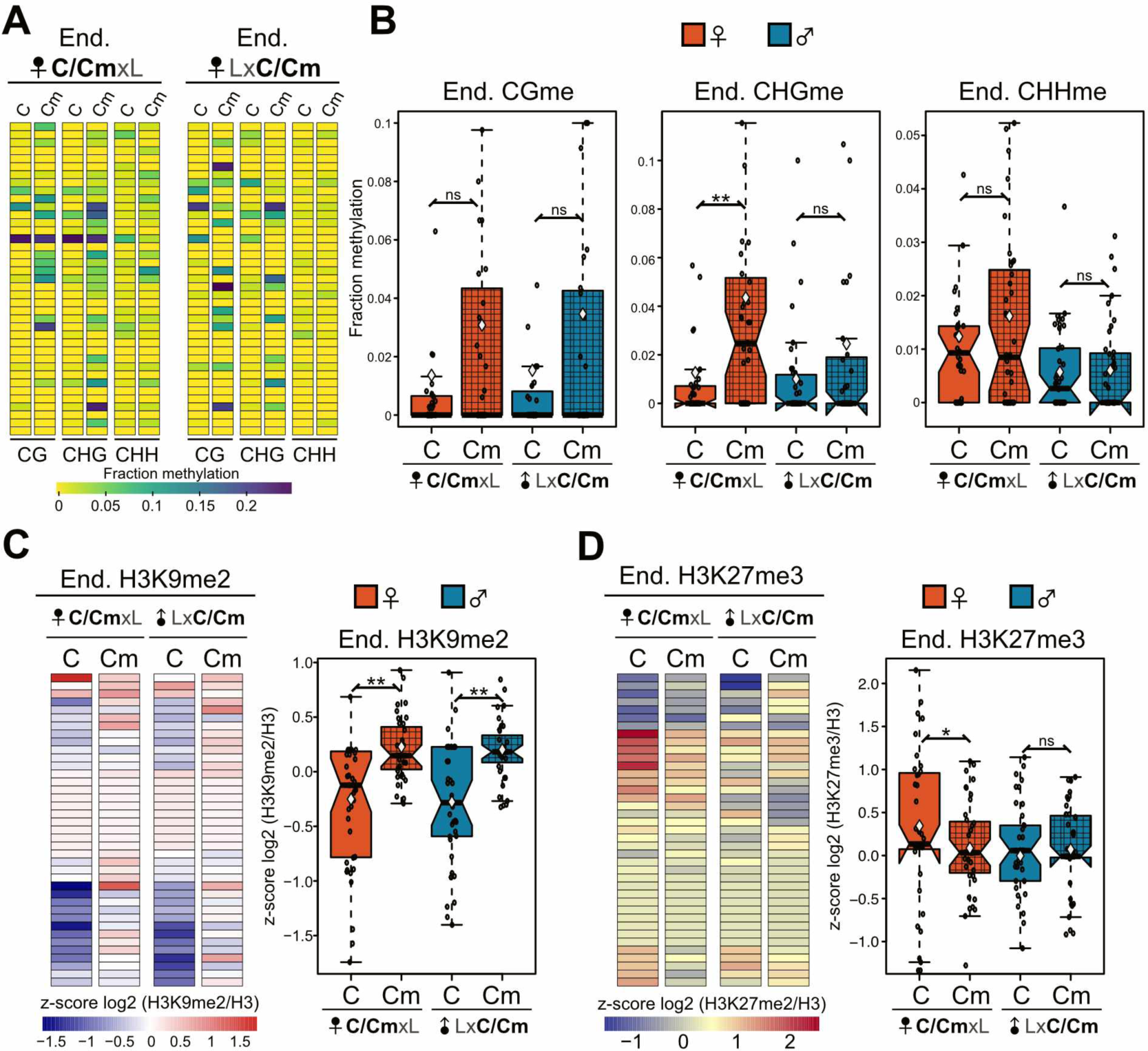
Parent-specific epigenetic effects in 100bp flanking regions of new *ONSEN* elements in TEmob. (**A**) Endosperm parental-specific DNA methylation in flanking regions of new *ONSEN* insertions in TEmob in comparison to WT. (**B**) Boxplot of the regions shown in the heat maps. Boxes show medians and the interquartile range (IQR), error bars show the full range excluding outliers. (**C**) Heatmaps (left) and boxplots (right) of endosperm parental- specific H3K9me2 in flanking regions of new *ONSEN* insertions in TEmob in comparison to WT. (**D**) Heatmaps (left) and boxplots (right) of endosperm parental-specific H3K27me3 in flanking regions of new *ONSEN* insertions in TEmob in comparison to WT. Smooth color boxes refer to crosses with WT, gridded pattern corresponds to crosses with TEmob. C, Col- 0 WT; Cm, Col-0 TEmob; L, L*er*. C/CmxL and LxC/Cm indicate crosses with either C or Cm. Wilcoxon test, \*\**P*-value <0.01; \**P*-value <0.05; ns, not significant.

In contrast to the observed enrichment of H3K9me2 at *ONSEN* flanking regions, we found that those regions were significantly depleted for H3K27me3 (**Figure 2D, Supplementary Figure S8B**), suggesting that *ONSEN* interrupts H3K27me3 deposition at the regions it inserts into. This decrease remained restricted to the flanking regions of new *ONSEN* insertions and did not impact H3K27me3 at the remaining parts of the genes (**Supplementary Figure S9B**). Depletion of H3K27me3 was restricted to the maternal genome, H3K27me3 levels were low at *ONSEN* insertion sites when paternally inherited and did not change before and after insertion (**Figure 2D**). These observations suggest that *ONSEN* insertion into H3K27me3 rich regions locally impairs H3K27me3 deposition, possibly due to recruitment and spreading of H3K9me2 (**Figure 2C and Supplementary Figure S9A**).

### Global changes of parental gene expression in TEmob endosperm

To understand whether *ONSEN* mobilization caused global expression changes in the endosperm, we generated parental-specific endosperm transcriptome profiles at 4DAP by reciprocally crossing TEmob Col (Cm) × L*er* (L) accessions (**Supplementary Table S7 and S8**). When TEmob was paternally inherited (LxCm), we observed a striking number of deregulated genes compared to WT (LxC), with 496 up- and 961 downregulated genes (FDR ≤ 0.01, LFC ±1.5) (**Figure 3A, B and Supplementary Table S9 and S10**). In contrast, in the reciprocal CmxL cross there were substantially fewer deregulated genes, with 74 up- and 239 downregulated genes compared to WT CxL (FDR ≤ 0.01, LFC ±1.5) (**Figure 3A, B and Supplementary Table S11 and S12**). 35 Transcription Factors (TFs) were among the upregulated genes when TEmob was the paternal parent (∼7% of the total upregulated genes), while only one TF was upregulated in the reciprocal CmxL cross (∼1% of the total upregulated) (**Figure 3C**, **Supplementary Table S13, S14 and Supplementary Figure S10**). These observations revealed a global parent-of-origin effect on gene expression in the TEmob endosperm, suggesting an epigenetic effect.

**Figure 3.**
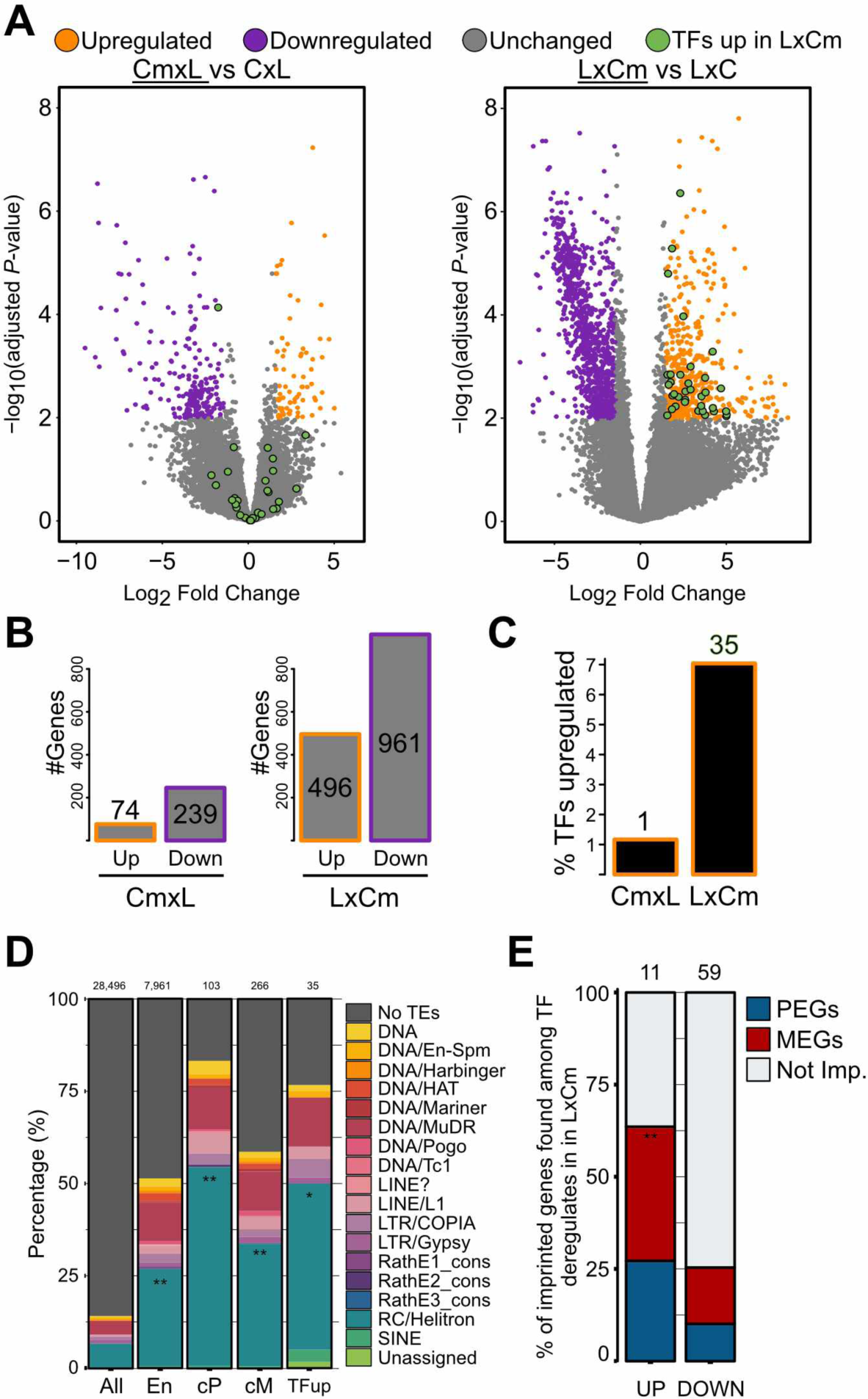
TEmob endosperm shows a parent-of-origin effect on differential gene expression in the endosperm. (**A**) Volcano plots showing differential gene expression of TEmob vs WT endosperm in reciprocal crosses with L*er* (FDR ≤ 0.01, LFC ±1.5). TFs, transcription factors. (**B**) Number of up- and downregulated genes in the indicated crosses. (**C**) Percentage of the deregulated genes annotated as Transcription Factors (TFs). (**D**) Percentage of transposable elements overlapping gene groups: All, All genes in TAIR genome; En, Genes expressed in Endosperm; cP, core PEGs and cM core MEGs are reported in at least two independent datasets; TFup: TFs upregulated in LxCm. Significance was tested using a Fisher test by comparing all groups to En (En was compared to All), \*\**P*-value <0.01; \**P*-value <0.05. (**E**) Percentage of imprinted TFs deregulated in LxCm. Not Imp., not imprinted genes; Hypergeometric test, \*\**P*-value <0.01; \**P*-value <0.05. C, Col-0 WT; Cm, Col-0 TEmob; L, L*er*.

To further understand the induced changes in gene expression by paternal TEmob, we analyzed the upregulated TFs in LxCm. Interestingly, of the 35 TF-encoding genes upregulated in LxCm endosperm, three (*AGL45*, *AGL48*, *AGL95*) are Type I MADS-box Mγ genes, with potential roles in endosperm development (**Supplementary Table S13**) (56, 57). We also observed that many of the upregulated TFs overlapped with TEs, particularly helitrons, resembling imprinted genes (**Figure 3D**). Of the upregulated TFs, 60% are imprinted genes with strong overrepresentation of MEGs (**Figure 3E**) (*P*-value<0.001, Hypergeometric Test). We thus conclude that paternal inheritance of TEmob results in upregulation of TFs in the endosperm, with many of them being MEGs, indicating a parent-of-origin epigenetic effect.

### Paternal hypomethylation in TEmob impacts MEGs in LxCm

Parent-of-origin transcriptional effects in the TEmob endosperm suggested global epigenetic changes characterizing the TEmob endosperm. Combined with heat shock, efficient *ONSEN* mobilization requires transient hypomethylation by zebularine, an inhibitor of DNA methylation (58) and α-amanitin, an inhibitor of RNA polymerase II (16). We therefore speculated that upregulation of MEG TFs is a consequence of hypomethylated and thus activated paternal alleles. To determine whether chemical hypomethylation impacted DNA methylation in TEmob four generations after treatment, we identified differentially methylated regions (DMRs) from vegetative aerial tissues. We identified 15, 055 hypomethylated and 5, 319 hypermethylated CG DMRs, revealing that the hypomethylated status was largely retained four generations after the chemically induced hypomethylation (**Figure 4A and B**). We identified only few non-CG DMRs in TEmob, yet there were about twice more CHG hypo- than hypermethylated regions (1, 395 compared to 717 DMRs, respectively). There were more hypermethylated CHH DMRs than hypomethylated CHH regions (1, 921 and 391, respectively), revealing that a subset of loci had undergone *de novo* CHH hypermethylation after treatment. Most of the CG DMRs overlapped with gene regions, 80% of DMRs were present in gene bodies and 10% in promoters, contrary to non-CG DMRs that were mostly localized in TEs (**Figure 4C**). We therefore conclude that TEmob maintains hypomethylation mainly in the CG context at least four generations after zebularine and α-amanitin treatment.

**Figure 4.**
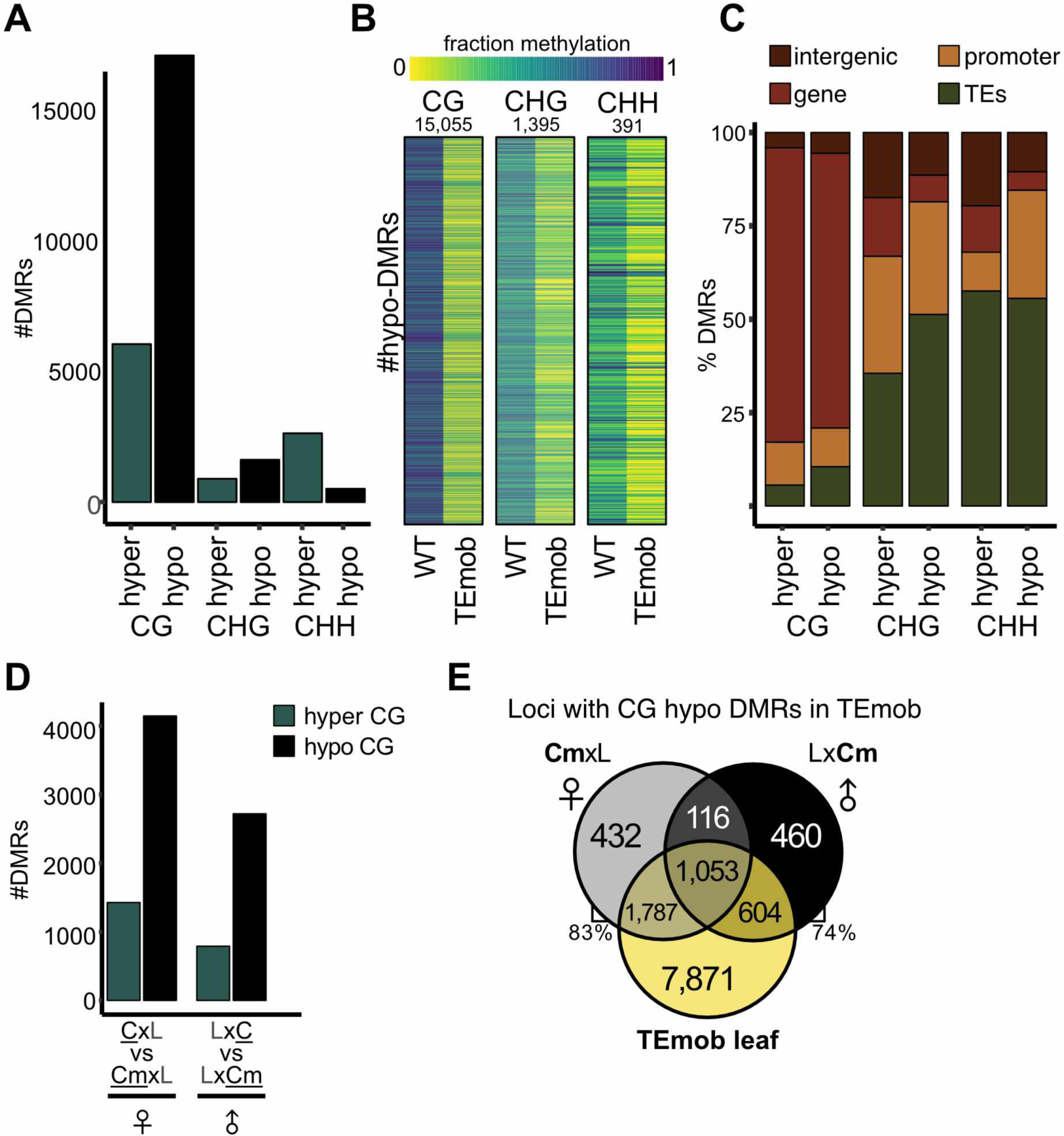
TEmob is highly hypo-methylated. (**A**) Differentially methylated regions (DMRs) in vegetative aerial tissues in TEmob. (**B**) Heat-maps of DMRs in all sequence contexts. (**C**) Annotation of TEmob DMRs in leaves. (**D**) TEmob DMRs detected in the endosperm by comparing reciprocal crosses of Col-L*er* (CxL) to Col TEmob-L*er* (CmxL). (**E**) Comparison of genes overlapping with TEmob DMRs detected in leaves and endosperm. C, Col-0 WT; Cm, Col-0 TEmob; L, L*er*. Parental-specific DMR comparisons are displayed in underscore.

To test how hypomethylation impacted TEmob endosperm, we identified parental-specific DMRs in the endosperm. Our analysis revealed that the TEmob genome transmitted hypo- and hypermethylated regions both as the maternal and paternal parent (**Figure 4D**), with most of the genes with endosperm DMRs overlapping DMRs found in leaves (**Figure 4E**). Out of a total of 4, 411 maternal hypomethylated DMRs in CmxL endosperm, 84% were CG DMRs, while the 2, 627 hypermethylated DMRs, were distributed over all sequence contexts (**Supplementary Figure S11A**). Similarly, in LxCm endosperm, 85% out of 2, 814 DMRs detected in the TEmob paternal genome were CG hypo DMRs, and 65% of those overlapped with maternal CG hypomethylated DMRs in CmxL endosperm. The 1, 008 hyper DMRs were distributed among CG, CHG and CHH contexts. Like in sporophytic tissues, parental-specific CG DMRs overlapped gene regions, whereas CHG and CHH DMRs were localized to TEs (**Supplementary Figure S11B**).

DNA hypomethylation could potentially lead to the activation of paternal alleles of MEGs as previously reported for *met1* (51, 59). Indeed, we observed a decreasing number of MEGs when TEmob was the paternal parent (**Figure 5A**). From the 437 reciprocal MEGs present in WT (31), 32.7% retained MEG expression only in the CmxL cross, and were biallelically expressed when TEmob was the paternal parent in LxCm endosperm (we will refer to those genes as Lost MEGs; LMEGs) (**Figure 5B, C**). Importantly, only 13% remained reciprocal MEGs in TEmob (we will refer to those genes as Strong MEGs; SMEGs) (**Figure 5B, C**). The remaining MEGs were reciprocally biallelic or showed a biased expression dependent on the accession. A stricter group of MEGs (MEGs found in at least two independent studies, referred to as “core” MEGs, cMEGs) confirmed these observations (**Figure 5B**). We concluded that most MEGs became biallelic when TEmob was the paternal parent. To test if paternal hypomethylation explained the reduced number of MEGs, we analyzed parental-specific DNAme in LMEGs. We however did not identify significant changes of DNA methylation in LMEGs in TEmob (**Supplementary Figure S12A**), indicating that paternal allele activation of MEGs did not globally connect with hypomethylation. In contrast to LMEGs, SMEGs showed a robust enrichment of CGme at the paternal alleles’ 5’ flanking regions in both WT and TEmob, suggesting a strong silencing of the paternal allele of these group of MEGs by CGme in the promoter regions (**Figure 5D, Supplementary Figure S12B**). Though paternal *met1* was shown to activate paternal MEG alleles, its effect is restricted to a handful of genes (51, 59).

**Figure 5.**
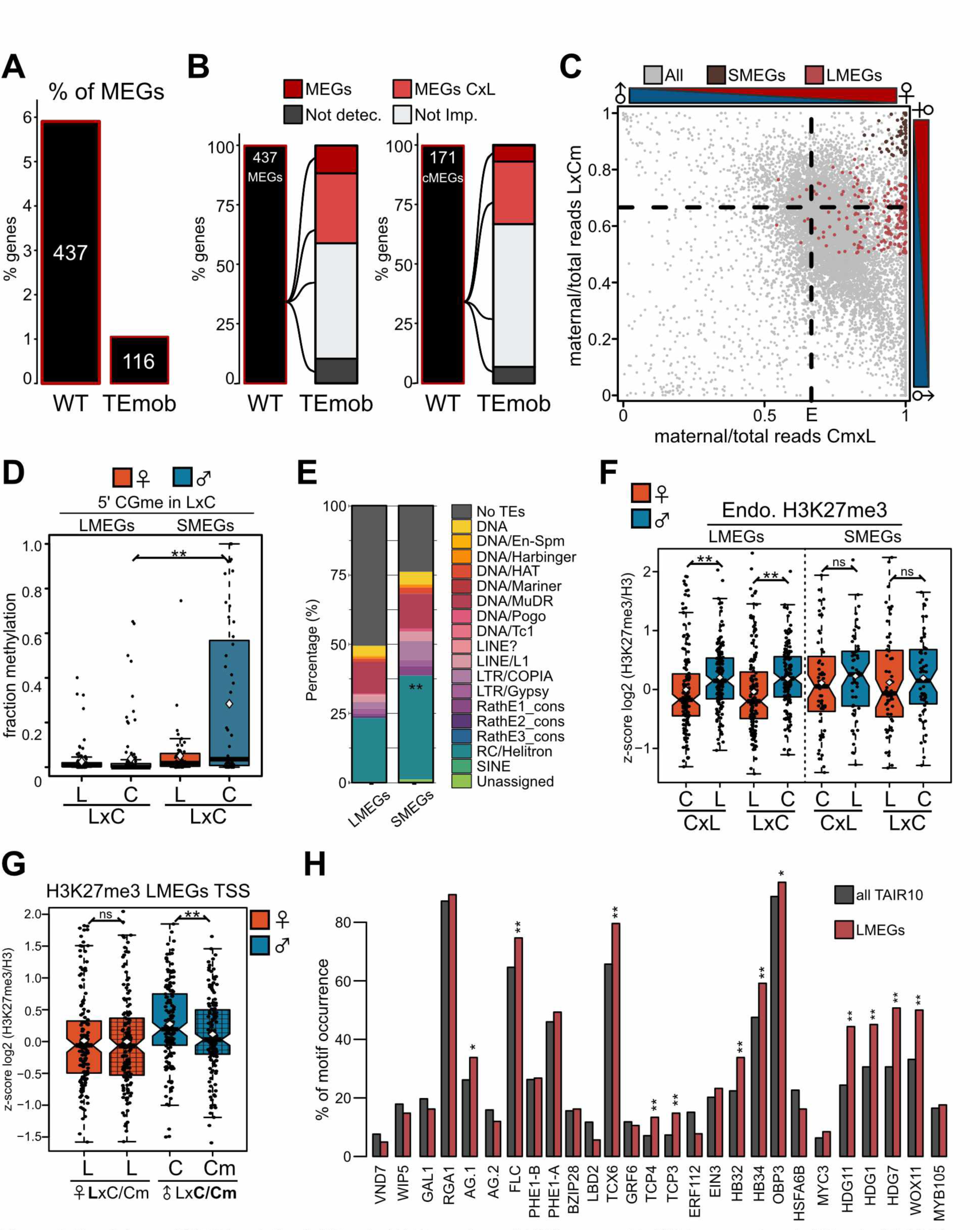
Breakdown of MEG imprinting in TEmob. (**A**) Percentage of MEGs present in WT in comparison to TEmob. (**B**) MEGs switch in TEmob. 437 MEGs in WT C-L dataset. (**C**) Scatter plot depicting parent-of-origin expression in TEmob comparing Lost MEGs (LMEGs) to Strong MEGs (SMEGs). (**D**) Boxplot showing CG methylation in upstream regions of LMEGs and SMEGs in LxC endosperm. (**E**) Percentage of transposable elements overlapping LMEGs and SMEGs. Fisher test comparing LMEGs vs SMEGs, \*\**P*-value <0.01. (**F**) Boxplots showing endosperm parental-specific H3K27me3 accumulation at LMEGs and SMEGs in WT endosperm. (**G**) Parental-specific H3K27me3 occupancy at the transcription start site region (TSS) in LMEGs in LxC TEmob (Cm) endosperm in comparison to WT (C). (**H**) Motif occurrence of upregulated transcription factors in LxCm endosperm in LMEGs and all genes annotated in TAIR10 (FIMO scan *P*-value =0.01). Fisher test, \*\**P*-value <0.01; \**P*-value <0.05. (D, F and G) Smooth color boxes refer to crosses with WT, gridded pattern refers to crosses with TEmob. C/CmxL and LxC/Cm indicate crosses with either C or Cm. Boxes show medians and the interquartile range (IQR), error bars show the full range excluding outliers. Wilcoxon test, \*\**P*-value <0.01; \**P*-value <0.05; ns, not significant. All MEGs, all MEGs in Col-L*er* dataset; cMEGs, core MEGs defined as those reported in at least two independent datasets; SMEGs, strong MEGs, MEGs did not change imprinting in TEmob paternal LxCm; LMEGs, Lost MEGs in TEmob paternal LxCm; Not detec., genes with no parental-specific data; Not Imp., not imprinted genes defined as genes not meeting criteria to be called as MEGs or PEGs or showing accession biased expression.

We tested if previously reported MEGs affected by *met1* were also affected in TEmob. Indeed 8 of the 16 MEGs losing imprinting upon *met1* pollination, were also affected in paternal TEmob endosperm (**Supplementary Table S15**). We therefore concluded that a limited number of LMEGs can be explained by hypomethylation in LxCm TEmob endosperm, while most MEGs lose their imprinted status because of methylation-independent changes.

### Ectopic expression of TFs potentially impairs MEG imprinting in TEmob

The paternal alleles of LMEGs were not marked by CGme, in contrast to the strongly methylated paternal alleles of SMEGs (**Figure 5D**). In addition, SMEGs showed a stronger association with TEs in their flanking regions than LMEGs, particularly with helitrons and LTR/Copia elements, suggesting a differential epigenetic regulation (**Figure 5E**). We thus tested whether paternal allele silencing of LMEGs may be mediated by repressive H3K27me3. We found that indeed LMEGs, in contrast to SMEGs, had significantly higher levels of H3K27me3 on the paternal compared to the maternal alleles, possibly accounting for paternal allele repression (**Figure 5F**). In LxCm TEmob there was a significant reduction of H3K27me3 particularly at regions close to the transcription start site (TSS) on the paternal alleles of LMEGs, which was not observed for SMEGs (**Figure 5G, Supplementary Figure S12C**). Based on these findings we speculated that increased TF activity (**Figure 3C**) may cause activation of the silenced paternal LMEG alleles and depletion of H3K27me3.

We identified 35 TFs to be upregulated in LxCm endosperm (**Figure 3C**). We thus wondered whether increased expression of TFs may explain the activation of silenced paternal MEG alleles. To test this idea, we identified binding motifs of the upregulated TFs and quantified their occurrence in the group of LMEGs (**Supplementary Table S16**). The predicted binding motifs were indeed present in LMEGs and several motifs were present in higher proportion than expected, including motifs for AGLs, HOMEODOMAIN GLABRA (HDG) and HOMEOBOX (HB) TF families as well as for TCX6 (**Figure 5H**). These results suggest that ectopic expression of TFs may contribute to the activation of the paternal allele of MEGs in the LxCm endosperm. One of the new *ONSEN* insertions occurred in an exon of *HDG3,* a transcription factor involved in endosperm cellularization (60). We found 14 commonly up- and 9 commonly downregulated genes in *hdg3-1* (60) and LxCm (**Supplementary Figure S13**). While the overlap with the upregulated genes was significant, the small number of commonly deregulated genes (**Supplementary Table S17**) make it unlikely that the insertion in *HDG3* accounts for the observed transcriptional changes in TEmob.

## DISCUSSION

Genomic imprinting has long been associated with the presence of TEs in the vicinity of genes; however, whether new TE insertions cause an instant effect on genomic imprinting has not been experimentally tested so far. As a proof of concept, we mobilized *ONSEN,* a heat shock responsive COPIA element and tested the consequences on genomic imprinting. We generated a line containing 40 new full-length homozygous *ONSEN* copies distributed across the five Arabidopsis chromosomes. The majority of the insertions were located in exons, confirming the insertion preference of *ONSEN* for coding genes (16). We further revealed a strong *ONSEN* insertion preference for imprinted genes, particularly MEGs. Less than 20% of the targeted MEGs contained a TE in the vicinity prior *ONSEN* insertion (**Supplementary Table S4 and S5**), revealing a weak association of MEGs and TEs. While the literature generally infers a strong association of TEs with imprinted genes, not all genes overlapping with TEs are imprinted, neither do all imprinted genes colocalize with a TE (13). Our observations indicate that the association of imprinted genes with TEs could also arise by the preferential insertion of TEs into imprinted genes, thus genomic imprinting may precede TE insertion. Determining the insertion preferences of TEs other than *ONSEN* will be important to test this hypothesis. In the case of *ONSEN*, the H2A.Z histone variant and H3K27me3 establish the chromatin landscape preferentially targeted by *ONSEN* (17). We found that MEGs are enriched for both, H2A.Z histone variant and H3K27me3, consistent with the preferential insertion of new *ONSEN* elements into MEGs.

Similar to previous observations for the remobilized *AtCOPIA93* element *Evadé* (*EVD*) (61), we found that new *ONSEN* insertions in TEmob were silenced in vegetative tissues, as evidenced by DNA methylation enrichment in all sequence contexts that extended into flanking regions. In the endosperm, new *ONSEN* elements were marked by H3K9me2 on both parental alleles, which co-occurred with CHGme on the maternal alleles. In contrast, there was only little CHHme on newly inserted *ONSEN* elements, consistent with low activity of the RdDM pathway in the early endosperm (10, 62). These data suggest that silencing of newly inserted *ONSEN* elements in the endosperm is mediated by H3K9me2 and partially independent of DNA methylation. Similarly, H3K9me2 deposition independent of DNA methylation was previously shown to occur in the Arabidopsis embryo, mediated by the histone methyltransferase SUVH9, whose activity is proposed to depend on small interfering RNAs (siRNAs) (55). *SUVH9* is a PEG (13) and recruits H3K9me2 in the endosperm (53); whether it has a role in silencing *ONSEN* in the endosperm remains to be shown.

We noticed a strong parent-of-origin effect on gene expression when TEmob was used as paternal parent (LxCm), with more than 32.7% of MEGs becoming biallelically expressed. We found that chemical TE mobilization caused and retained hypomethylation in TEmob four generations after treatment, extending previous findings showing stable inheritance of hypomethylated regions for two generations (63). We therefore hypothesized that the activation of paternal MEG alleles in LxCm is a consequence of paternal allele hypomethylation, explaining the parent-of-origin effect on gene deregulation. However, DNA methylation analysis failed to globally connect hypomethylation with the loss of MEG imprinting. Using TEmob as paternal parent we noticed a strong upregulation of TFs (**Figure 3C**). Interestingly, we found that biallelically expressed MEGs (LMEGs) contained binding motifs for the upregulated TFs (**Figure 5G, Supplementary Table S16**). While MEGs that remained imprinted were highly enriched for CGme in the promoter region of paternal alleles, the paternal alleles of LMEGs were not marked by CGme. Instead, they had significantly higher levels of H3K27me3 on paternal compared to the maternal alleles, which were significantly reduced at their TSS region in TEmob. These observations suggest that ectopic expression of TFs caused activation of the silenced paternal LMEG alleles by depletion of H3K27me3. In crosses using *met1* as paternal parent (51, 59), only few (16) MEGs lose their imprinting status (**Supplementary Table S15**). One possible explanation for the large number of MEGs affected in TEmob is that loss of MET1 function mainly affects CG methylation, while zebularine impairs all DNA methylation pathways, with predictably larger effects (64). Together, we conclude that the observed global breakdown of MEG imprinting by paternal TEmob is largely attributed to upregulated TFs that activate the silenced paternal alleles of MEGs.

## DATA AVAILABILITY

Original data files for WGS, WSBS, RNAseq and CUT&Tag sequencing can be obtained from the NCBI Gene Expression Omnibus (GSE260822). The assembled TEmob genome sequence can be obtained from the ENA (PRJEB73629).

## SUPPLEMENTARY DATA

Supplementary Figures

**Supplementary Figure S1.** Establishment of TEmob line. Transposable element mobilization and doubled haploid induction. alpha-amanitin (A); zebularine (Z); Heat Shock (HS); Doubled-Haploid (DH).

**Supplementary Figure S2.** New *ONSEN* in TEmob. (**A**) Graphical representation of the chromosomal distribution of all 40 *AtONSEN_mob* in TEmob (**B**) PCR amplification of randomly selected *AtONSEN_mob* from Col-TEmob and WT Col-0 genomic DNA using a primer in *ONSEN* in combination with a primer located at a new flanking region. Primers indicated in Supplementary Table S1. *OP*, ONSEN primer; *FP*, Flanking primer.

**Supplementary Figure S3.** Phylogenetic tree based on DNA sequence alignments of new *ONSEN* in TEmob and native full-length *ONSEN* in Col-0. Underscored names indicate the most related native *ONSEN* to new copies in TEmob.

**Supplementary Figure S4.** DNA methylation of native *ATCOPIA78* in aerial vegetative tissues. (**A**) Heatmap showing DNA methylation in all sequence contexts of individual *ATCOPIA78* family members in leaf tissues in WT and TEmob. Mobile *ONSEN* copies are underlined in red. Black asterisks, full-length *ATCOPIA78* in Col-0; Red asterisk, mobile *ONSEN* copies reported in the literature; Blue asterisk, reported with high extrachromosomal circular DNA abundance (eccDNA) in Roquis *et al,* 2021. (**B**) Bulk DNA methylation as in **A**. Wilcoxon test, ns, not significant.

**Supplementary Figure S5.** Accumulation of H2A.Z and H3K27me3 in leaf tissues. (**A**) H2A.Z and (**B**) H3K27me3 accumulation over *ONSEN* targeted genes (*ONSEN* TG) in TEmob. . Comparisons against random samples (RS) of the same number of genes. Wilcoxon test, \*\**P*- value <0.01; \**P*-value <0.05; ns, not significant.

**Supplementary Figure S6.** Parental-specific endosperm DNA methylation in all sequence contexts in (**A**). cMEGs, (**B**) cPEGs, and (**C**) Biallelic genes in reciprocal crosses with Col-0 (C) or TEmob (Cm) and L*er* (L). C, Col-0 WT; Cm, Col-0 TEmob; GB, Gene Body. Core MEGs and PEGs are defined as imprinted in at least two independent studies. Smooth color boxes refer to crosses with WT, gridded pattern refers to crosses with TEmob. C/CmxL and LxC/Cm indicate crosses with either C or Cm. \*\**P*-value <0.01; \**P*-value <0.05; ns, not significant.

**Supplementary Figure S7.** Parental-specific accumulation of H3K27me3 (**A, B**) and H3K9me2 (**C, D**) in core MEGs (cMEGs, **A, C**) and core PEGs (cPEGs, **B, D**) in WT reciprocal CxL and LxC endosperm. C, Col-0; L, L*er*; RS, random sample. Comparisons against random samples of the same number of genes. Wilcoxon test, \*\**P*-value <0.01; \**P*-value <0.05; ns, not significant.

**Supplementary Figure S8.** Genome Browser Screenshots of new *ONSEN* insertions in TEmob and WT. (**A**) Parental-specific H3K9m2 in the endosperm. (**B**) Parental-specific H3K27me3 in the endosperm. Blue triangles show the location of newly inserted *ONSEN* in TEmob. C, Col-0 WT; Cm, Col-0 TEmob; L, L*er*.

**Supplementary Figure S9.** Parental-specific H3K9me2 (**A**) and H3K27me3 (**B**) accumulation over genes targeted by *ONSEN* in TEmob and WT endosperm (C/CmxL and LxC/Cm). C, Col- 0 WT; Cm, Col TEmob; L, L*er*; RS, Random Sample. Smooth color boxes refer to crosses with WT, gridded pattern refers to crosses with TEmob. C/CmxL and LxC/Cm indicate crosses with either C or Cm. Wilcoxon test, \*\**P*-value <0.01; \**P*-value <0.05; ns, not significant.

**Supplementary Figure S10.** Heat map showing expression of the transcription factors upregulated in LxCm compared to WT LxC. r, replicate; C, Col-0 WT; Cm, Col TEmob; L, L*er*.

**Supplementary Figure S11.** Parental-specific differentially methylated regions (DMRs) in the TEmob endosperm. (**A**) Parental specific CGme, CHG and CHH DMRs in the endosperm. **(B**) Annotation of parental-specific DMRs in the endosperm involving TEmob (Cm). C, Col-0 WT; Cm, Col TEmob; L, L*er*. Parental-specific DMR comparisons are visualized by underscore.

**Supplementary Figure S12.** Parental-specific DNA methylation in Lost MEGs (LMEGs) (**A**) and strong MEGs (SMEGs) (**B**) in LxC/Cm endosperm at 5’ UTR (left) and gene body (GB) (right). (**C**) Parental-specific accumulation of H3K27me3 at the transcriptional start site (TSS) of SMEGs in LxC/Cm endosperm. (**D**) Parental-specific accumulation of H3K9me2 in LMEGs and SMEGs n LxC/Cm endosperm. Smooth color boxes refer to crosses with WT, gridded pattern refers to crosses with TEmob. C, Col-0 WT; Cm, Col-0 TEmob; L, L*er*. C/CmxL and LxC/Cm indicate crosses with either C or Cm. Wilcoxon test, \*\**P*-value <0.01; \**P*-value <0.05; ns, not significant.

**Supplementary Figure S13.** Comparison of transcriptome differences in *hdg3-1* and TEmob. (**A**)Volcano plot showing upregulated and downregulated genes in TEmob LxCm endosperm and *hdg3-1* (Pignatta *et al* 2018). (**B**) Overlap of upregulated (top) and downregulated (bottom) genes in TEmob LxCm endosperm and *hdg3-1*. Hypergeometric test of significance. \*\**P*-value <0.01; ns, not significant.

Supplementary Tables File 1

**Supplementary Table S1.** List of primers.

**Supplementary Table S2.** Summary of new *ONSEN* insertions in TEmob.

**Supplementary Table S3.** Compilation of reported imprinted genes for Arabidopsis.

**Supplementary Table S4.** Parent-of-origin expression of *ONSEN* targeted genes prior to *ONSEN* insertion (1kb up- or downstream of ORFs).

**Supplementary Table S5.** Parent-of-origin expression of *ONSEN* targeted genes or genes in the vicinity of novel *ONSEN* insertions in the hcLines of Roquis et al 2021 (1kb up- or downstream ORFs).

**Supplementary Table S6.** Parent-of-origin expression of *ONSEN* targeted genes.

File 2

**Supplementary Table S7**. Full table with TPM (transcript per million) and CPM (counts per million) for all the RNAseq replicates used for differential gene expression analysis.

File 3

**Supplementary Table S8.** Parent-of-origin expression in reciprocal crosses of TEmob (Cm): CmxL; LxCm at 4DAP.

File 4

**Supplementary Table S9.** Full table with Differential Gene Expression analysis of LxCm (TEmob) compared to LxC (WT) endosperm and overlap with DMRs in leaves and endosperm together with parent-of-origin expression in WT and TEmob endosperm.

**Supplementary Table S10.** Genes up- or downregulated in LxCm (TEmob) compared to LxC (WT). DMRs present in Cm (TEmob) in the endosperm are indicated.

File 5

**Supplementary Table S11.** Full table with differential gene expression data on CmxL (TEmob) compared to CxL (WT) endosperm and overlap with DMRs in leaves and endosperm together with parent-of-origin expression data.

**Supplementary Table S12.** Genes up- or downregulated in CmxL (TEmob) compared to CxL (WT). DMRs present in Cm (TEmob) in endosperm and leaves are indicated.

File 6

**Supplementary Table S13.** Transcription factors up- or downregulated in LxCm (TEmob) compared to LxC (WT). DMRs present in Cm (TEmob) in endosperm and leaves are indicated. Overlap with TEs 1kb up- or downstream of ORFs or within introns is indicated.

**Supplementary Table S14.** Transcription factors up- or downregulated in CmxL (TEmob) compared to CxL (WT). DMRs present in Cm (TEmob) in the endosperm and leaves are indicated.

**Supplementary Table S15.** Parent-of-origin expression of MEGs reported to be affected by paternal *met1* in TEmob endosperm.

**Supplementary Table S16.** Predicted motifs for transcription factors upregulated in LxCm (TEmob).

**Supplementary Table S17**. Common upregulated and downregulated genes in LXCm (TEmob) endosperm and *hdg3-1*.

## AUTHOR CONTRIBUTIONS

*Author contributions*: CK and GDTDL conceptualized the manuscript; JB generated *ONSEN* mobilization lines. WBJ and KS assembled the TEmob genome; GDTDL performed the experiments; GDTDL, CK and JS analyzed the data; GDTDL and CK wrote the manuscript. All authors read and commented on the manuscript.

## ACKNOWLEDGEMENTS

### FUNDING

This work was supported by Knut and Alice Wallenberg Foundation grant 2018-0206 (CK), Knut and Alice Wallenberg Foundation grant 2019-0062 (CK), and the Max Planck Society, Germany.

