## Supplemental Figures for "Epigenetic and transcriptional consequences of chemically induced transposon mobilization in the endosperm"

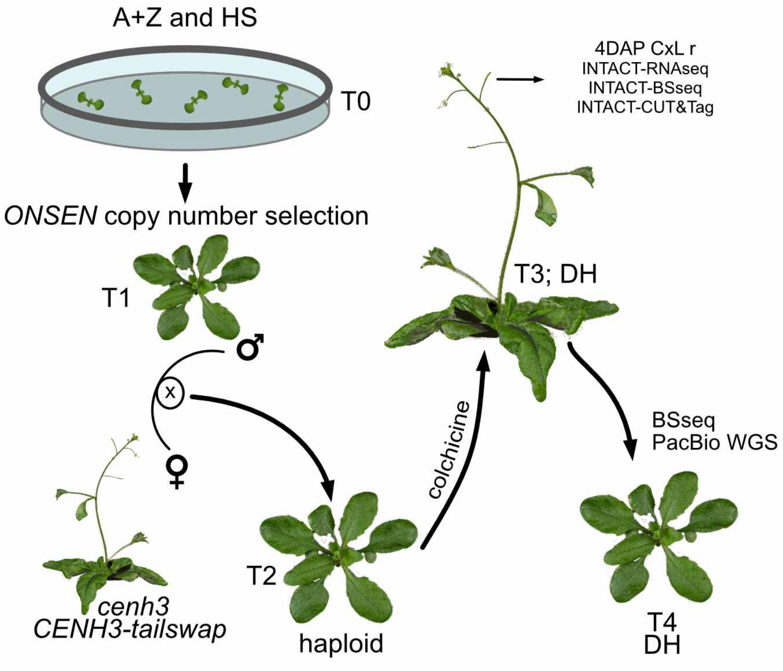


**Supplementary Figure S1.** Establishment of TEmob line. Transposable element mobilization and doubled haploid induction. alpha-amanitin (A); zebularine (Z); Heat Shock (HS); Doubled-Haploid (DH).

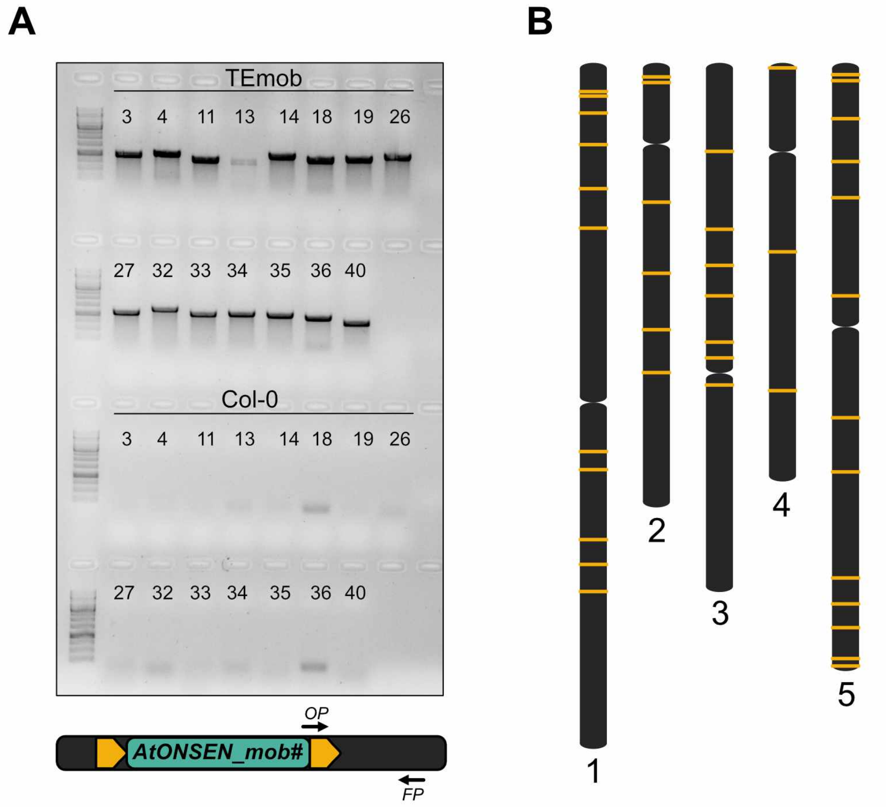


**Supplementary Figure S2.** New *ONSEN* in TEmob. (**A**) Graphical representation of the chromosomal distribution of all 40 *AtONSEN_mob* in TEmob (**B**) PCR amplification of randomly selected *AtONSEN_mob* from Col-TEmob and WT Col-0 genomic DNA using a primer in *ONSEN* in combination with a primer located at a new flanking region. Primers indicated in Supplementary Table S1. *OP*, ONSEN primer; *FP*, Flanking primer.

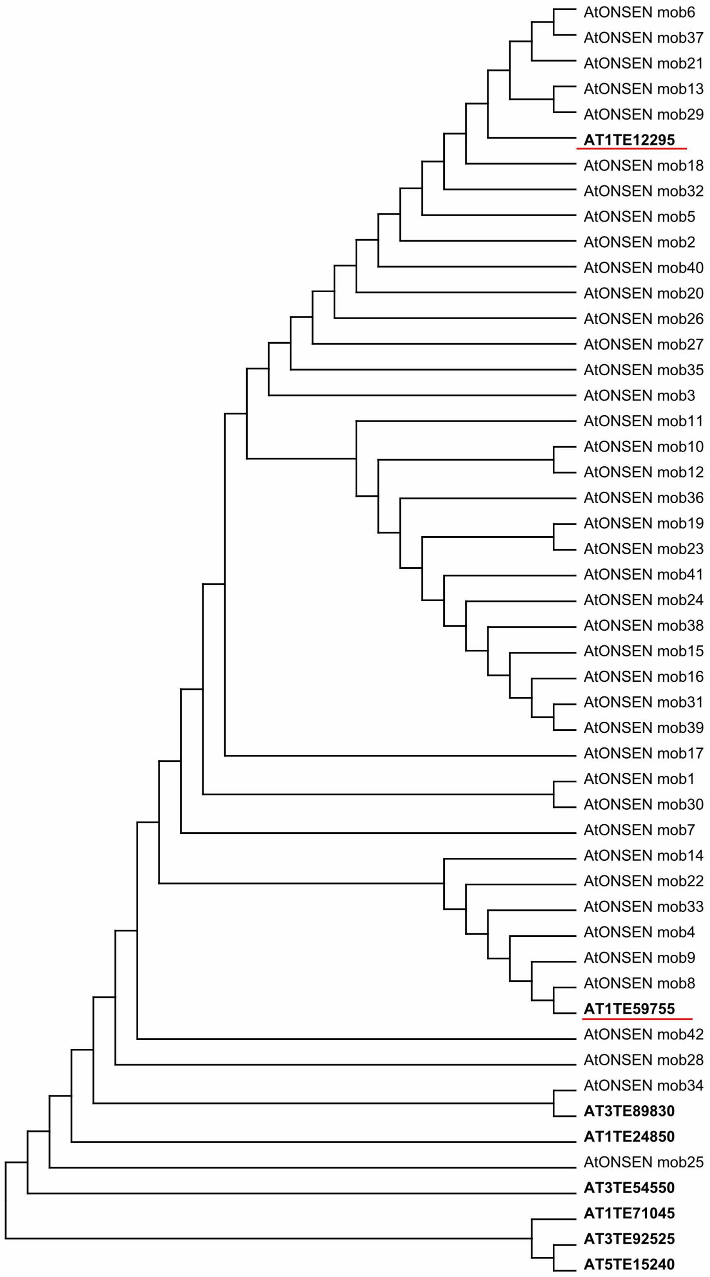


**Supplementary Figure S3.** Phylogenetic tree based on DNA sequence alignments of new *ONSEN* in TEmob and native full-length *ONSEN* in Col-0. Underscored names indicate the most related native *ONSEN* to new copies in TEmob.

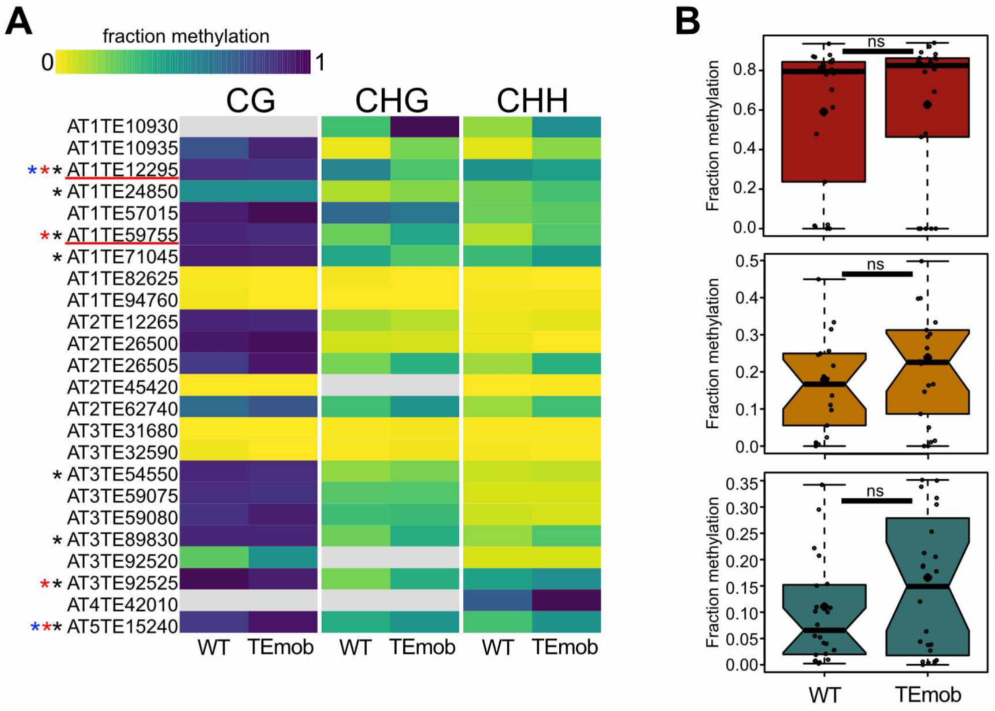


**Supplementary Figure S4.** DNA methylation of native *ATCOPIA78* in aerial vegetative tissues. (**A**) Heatmap showing DNA methylation in all sequence contexts of individual *ATCOPIA78* family members in leaf tissues in WT and TEmob. Mobile *ONSEN* copies are underlined in red. Black asterisks, full-length *ATCOPIA78* in Col-0; Red asterisk, mobile *ONSEN* copies reported in the literature; Blue asterisk, reported with high extrachromosomal circular DNA abundance (eccDNA) in Roquis *et al,* 2021. (**B**) Bulk DNA methylation as in **A**. Wilcoxon test, ns, not significant.

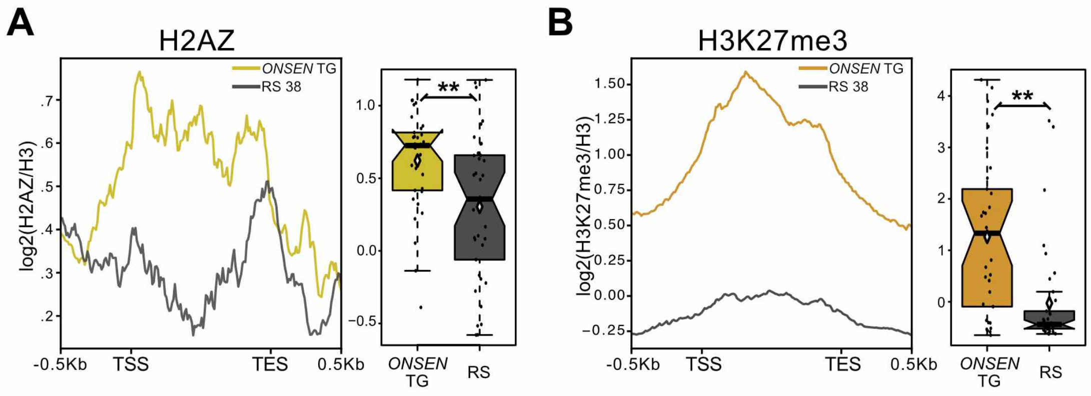


**Supplementary Figure S5.** Accumulation of H2A.Z and H3K27me3 in leaf tissues. (**A**) H2A.Z and (**B**) H3K27me3 accumulation over *ONSEN* targeted genes (*ONSEN* TG) in TEmob. . Comparisons against random samples (RS) of the same number of genes. Wilcoxon test, ***P*-value <0.01; **P*-value <0.05; ns, not significant.

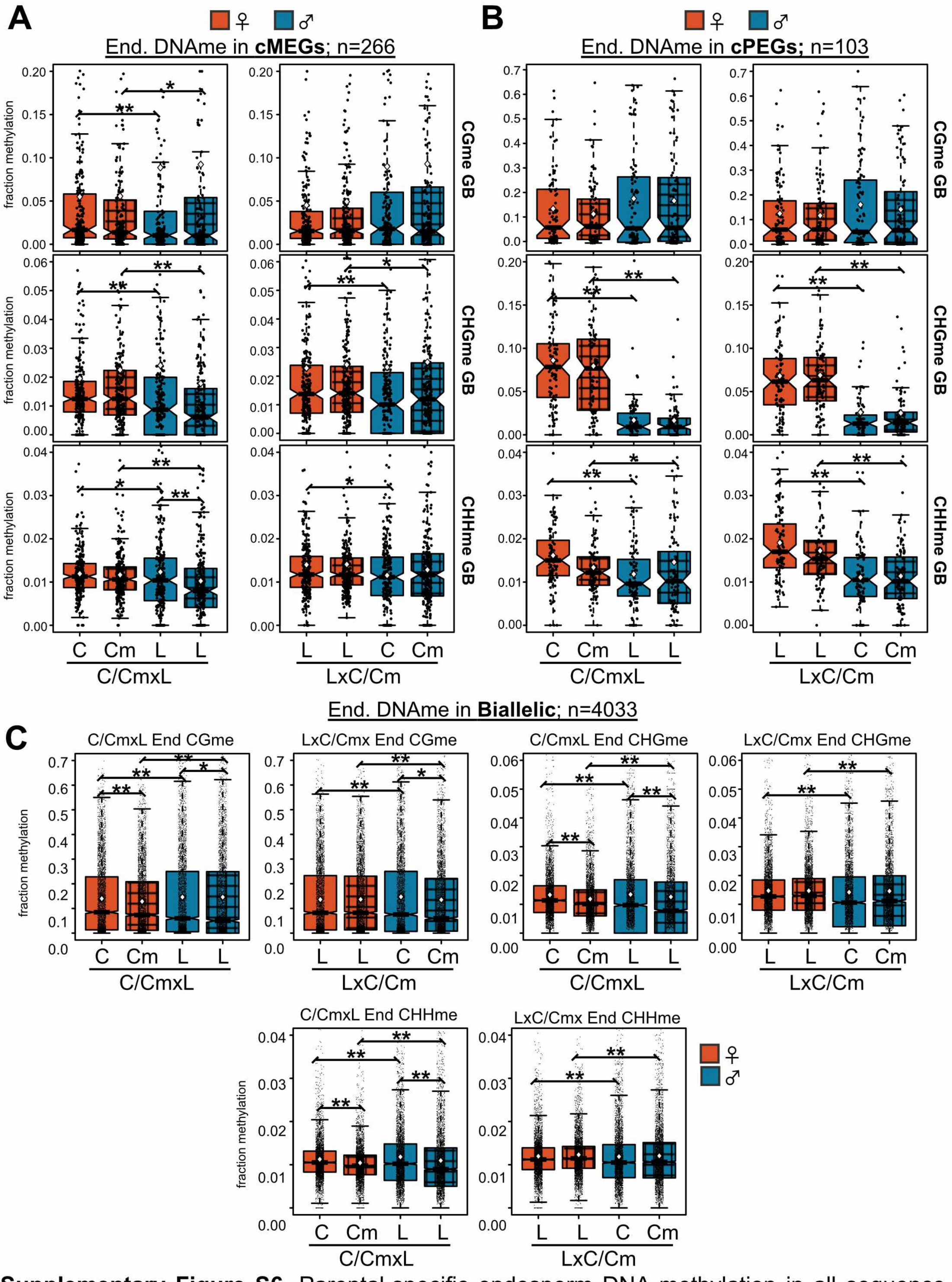


**Supplementary Figure S6.** Parental-specific endosperm DNA methylation in all sequence contexts in (**A**). cMEGs, (**B**) cPEGs, and (**C**) Biallelic genes in reciprocal crosses with Col-0 (C) or TEmob (Cm) and L*er* (L). C, Col-0 WT; Cm, Col-0 TEmob; GB, Gene Body. Core MEGs and PEGs are defined as imprinted in at least two independent studies. Smooth color boxes refer to crosses with WT, gridded pattern refers to crosses with TEmob. C/CmxL and LxC/Cm indicate crosses with either C or Cm. ***P*-value <0.01; **P*-value <0.05; ns, not significant.

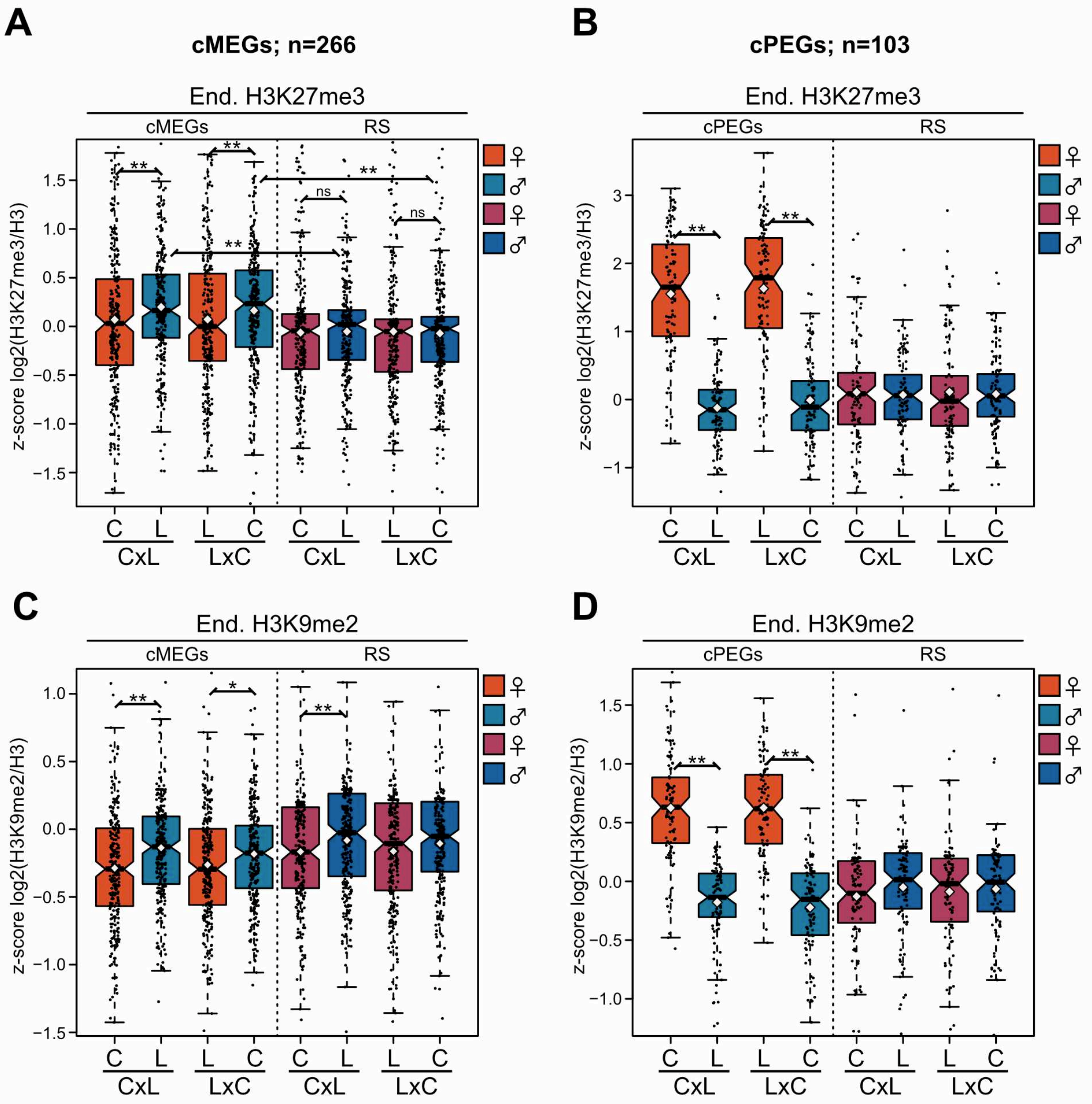


**Supplementary Figure S7.** Parental-specific accumulation of H3K27me3 (**A,B**) and H3K9me2 (**C,D**) in core MEGs (cMEGs, **A,C**) and core PEGs (cPEGs, **B,D**) in WT reciprocal CxL and LxC endosperm. C, Col-0; L, L*er*; RS, random sample. Comparisons against random samples of the same number of genes. Wilcoxon test, ***P*-value <0.01; **P*-value <0.05; ns, not significant.

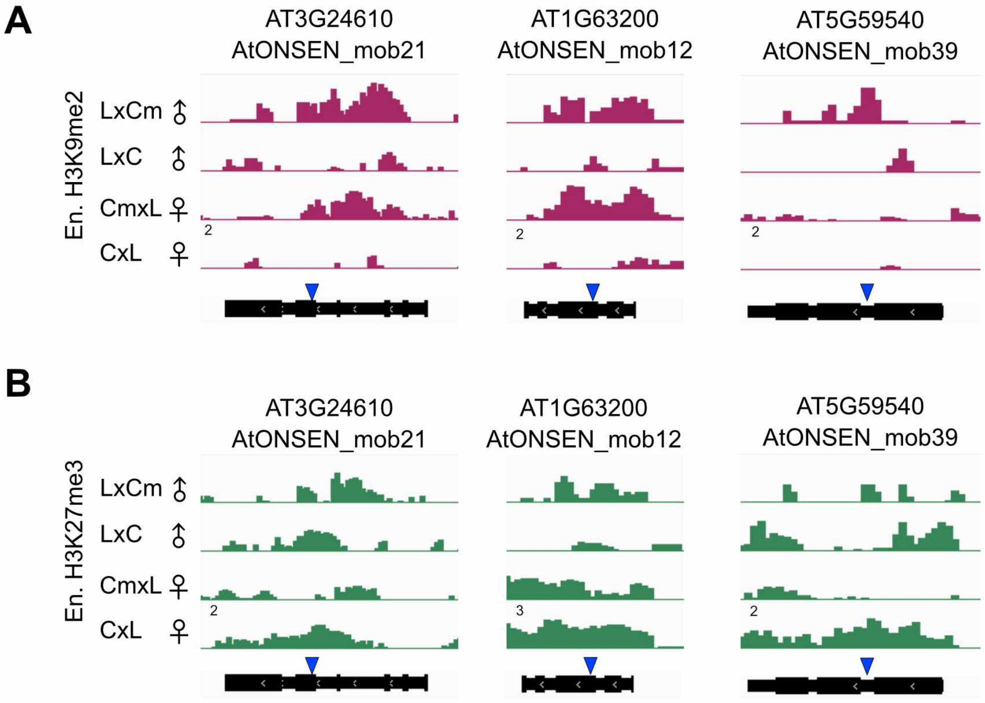


**Supplementary Figure S8.** Genome Browser Screenshots of new *ONSEN* insertions in TEmob and WT. (**A**) Parental-specific H3K9m2 in the endosperm. (**B**) Parental-specific H3K27me3 in the endosperm. Blue triangles show the location of newly inserted *ONSEN* in TEmob. C, Col-0 WT; Cm, Col-0 TEmob; L, L*er*.

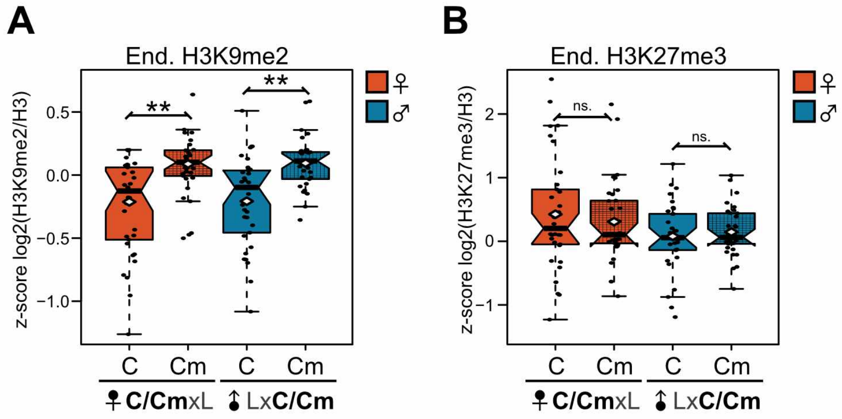


**Supplementary Figure S9.** Parental-specific H3K9me2 (**A**) and H3K27me3 (**B**) accumulation over genes targeted by *ONSEN* in TEmob and WT endosperm (C/CmxL and LxC/Cm). C, Col-0 WT; Cm, Col TEmob; L, L*er*; RS, Random Sample. Smooth color boxes refer to crosses with WT, gridded pattern refers to crosses with TEmob. C/CmxL and LxC/Cm indicate crosses with either C or Cm. Wilcoxon test, ***P*-value <0.01; **P*-value <0.05; ns, not significant.

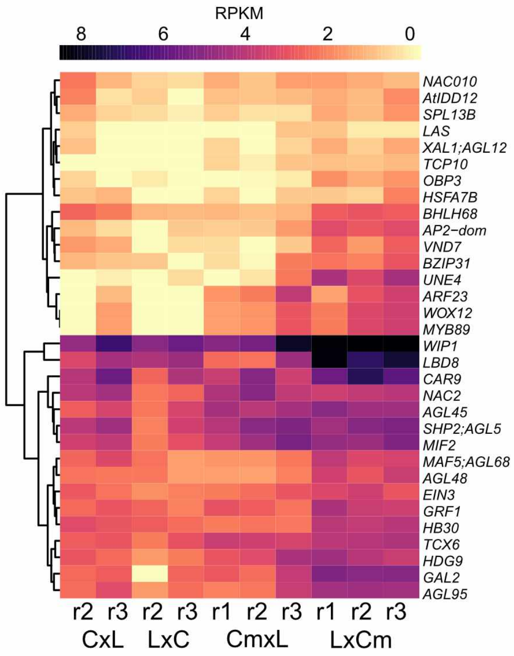


**Supplementary Figure S10.** Heat map showing expression of the transcription factors upregulated in LxCm compared to WT LxC. r, replicate; C, Col-0 WT; Cm, Col TEmob; L, L*er*.

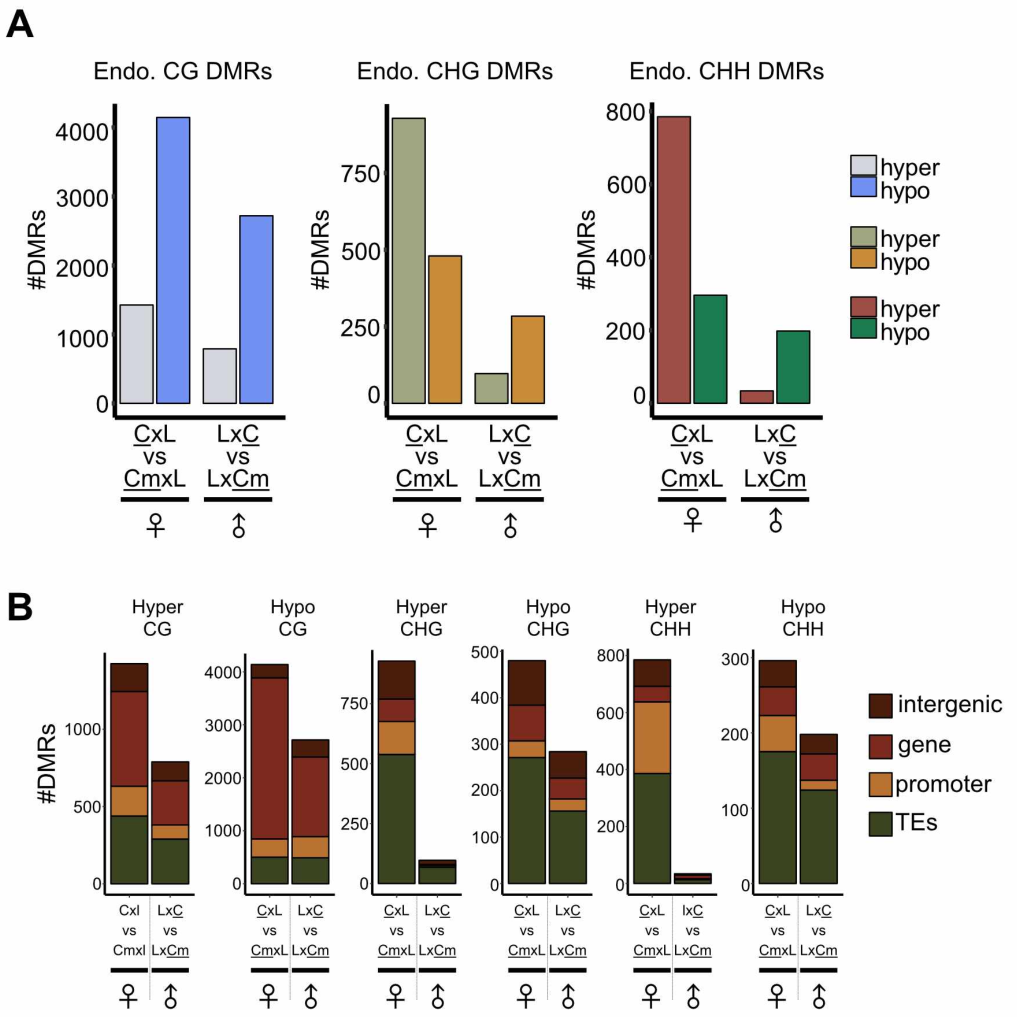


**Supplementary Figure S11.** Parental-specific differentially methylated regions (DMRs) in the TEmob endosperm. (**A**) Parental specific CGme, CHG and CHH DMRs in the endosperm. **(B)** Annotation of parental-specific DMRs in the endosperm involving TEmob (Cm). C, Col-0 WT; Cm, Col TEmob; L, L*er*. Parental-specific DMR comparisons are visualized by underscore.

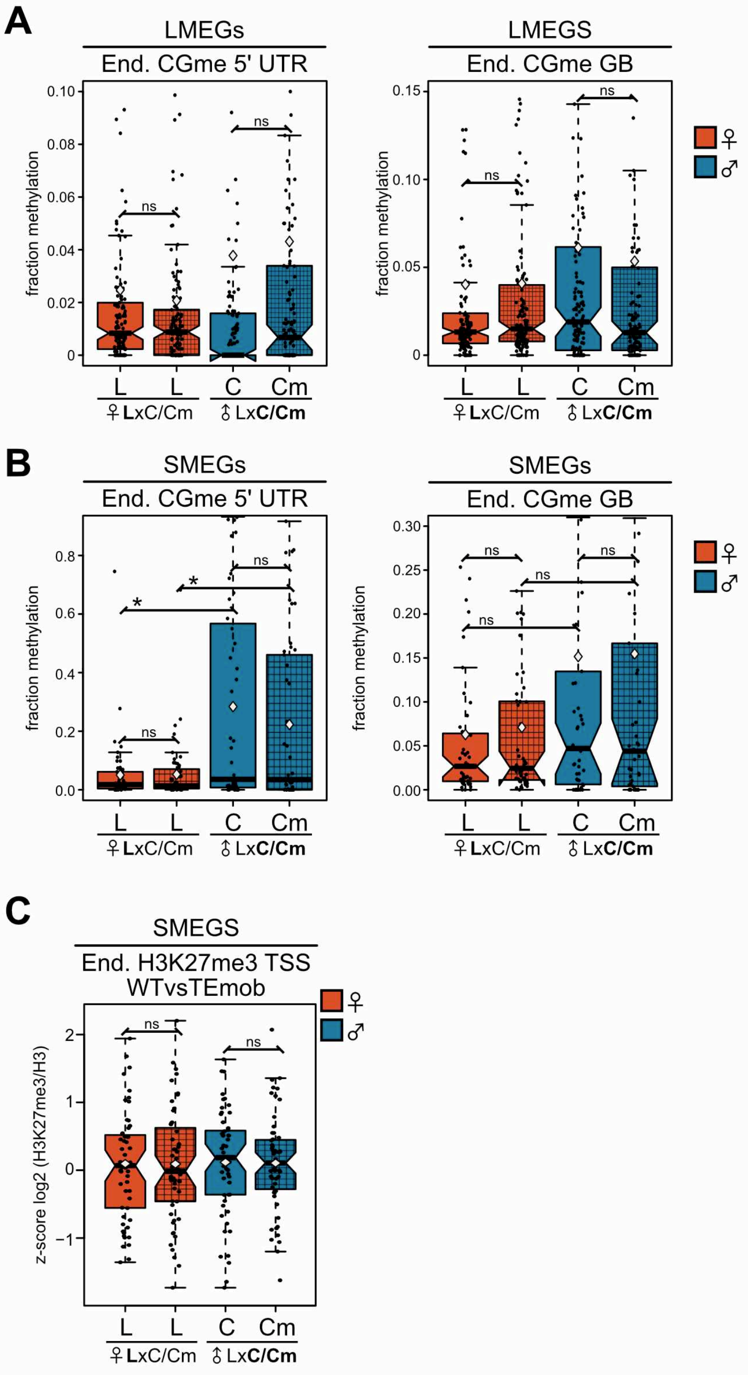


**Supplementary Figure S12.** Parental-specific DNA methylation in Lost MEGs (LMEGs) (**A**) and strong MEGs (SMEGs) (**B**) in LxC/Cm endosperm at 5' UTR (left) and gene body (GB) (right). (**C**) Parental-specific accumulation of H3K27me3 at the transcriptional start site (TSS) of SMEGs in LxC/Cm endosperm. (**D**) Parental-specific accumulation of H3K9me2 in LMEGs and SMEGs n LxC/Cm endosperm. Smooth color boxes refer to crosses with WT, gridded pattern refers to crosses with TEmob. C, Col-0 WT; Cm, Col-0 TEmob; L, L*er*. C/CmxL and LxC/Cm indicate crosses with either C or Cm. Wilcoxon test, ***P*-value <0.01; **P*-value <0.05; ns, not significant.

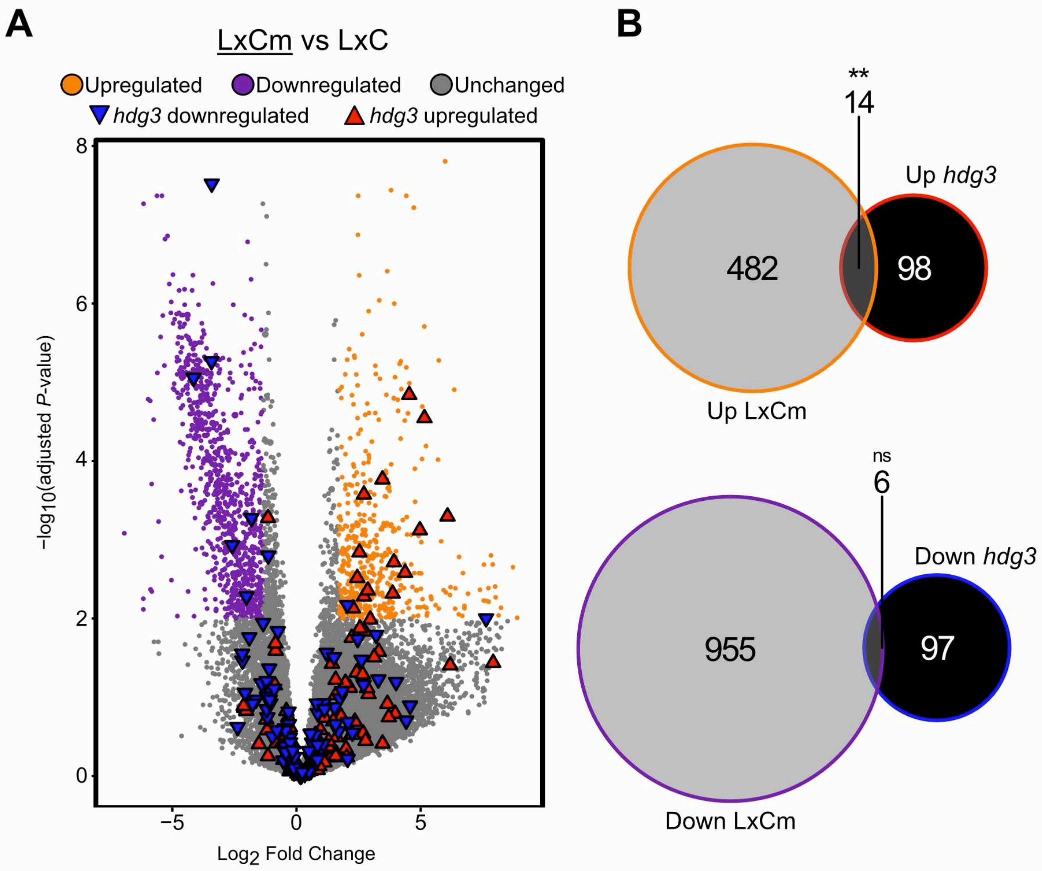


**Supplementary Figure S13.** Comparison of transcriptome differences in *hdg3-1* and TEmob. (**A**)Volcano plot showing upregulated and downregulated genes in TEmob LxCm endosperm and *hdg3-1* (Pignatta *et al* 2018). (**B**) Overlap of upregulated (top) and downregulated (bottom) genes in TEmob LxCm endosperm and *hdg3-1*. Hypergeometric test of significance. ***P*-value <0.01; ns, not significant.
